# Image-grounded encoding models reveal distinct temporal profiles of naturalistic object and scene processing in the human brain

**DOI:** 10.64898/2026.04.24.720581

**Authors:** Niklas Müller, H. Steven Scholte, Iris I. A. Groen

## Abstract

In real-world vision, the human brain needs to process large amounts of information to effectively interact with its environment. It is well established that our visual system has specialized regions to process information efficiently, such as scene-, face-, and object-selective areas, which can be uncovered using functional magnetic resonance imaging (fMRI). However, functional localizers typically use artificially separated stimuli (e.g. scene backgrounds vs. object cutouts), and fMRI signals are insensitive to potential temporal processing differences. Here, we identify temporal signatures of object and scene processing in real-world visual environments by building image-grounded brain-predictive encoding models of human electroencephalography (EEG) responses. In a large set of high-resolution natural images, we first separate object-bound from scene-bound features on a per-image basis, and then used these features to train separate deep neural network-based encoding models to predict EEG responses to the full, intact images. We find that encoding models trained on object-bound features consistently exhibit a delayed temporal encoding profile compared to models trained on scene-bound features. Control analyses confirm the robustness of this delayed object encoding, showing that a consistent separation of objects from scene parts is needed to achieve a temporal distinction. Using these distinct encoding profiles as processing templates, we reveal the typicality of individual image elements as objects or part of the scene. These findings demonstrate that object and scene processing in the human brain unfold differentially over time and indicate that image-grounded encoding models are helpful tools for isolating components of naturalistic visual perception.

## Introduction

In everyday vision, the human brain rapidly and efficiently processes vast amounts of visual information from real-world environments. This information forms the basis for a wide range of perceptual decisions, for example how to navigate the environment (Bonner and Epstein, 2017; Bartnik et al., 2025b), or how to use and manipulate objects (Bracci and Peelen, 2013; Baldassano et al., 2017). To achieve this feat, the visual system is thought to process visual inputs in a parallelized manner (Ballard, 1986; Rousselet et al., 2004; Nassi and Callaway, 2009), dedicating distinct neural pathways and cortical areas to processing specific types of visual information (Zeki et al., 1991). For example, visual cortex contains distinct regions involved in processing scene-specific information Epstein and Kanwisher (1998); Epstein and Baker (2019); Bartnik and Groen (2023) versus object-specific information (Kourtzi and Kanwisher, 2001; Riesenhuber and Poggio, 2002; Grill-Spector et al., 2001).

Although this functional specialization is well established (Kanwisher, 2010), two gaps remain in our understanding of object- and scene-specific visual processing in the brain. First, their temporal processing trajectories remain poorly understood. Object- and scene-specific components have been identified in electroencephalography (EEG) recordings (Harel et al., 2016), and multivariate decoding studies have elucidated stages of processing during either object-(Simanova et al., 2010; Cichy et al., 2013; Grootswagers et al., 2019) or scene-(Cichy and Teng, 2017; Lowe et al., 2018; Ramkumar et al., 2016) perception, but most prior research has focused on spatial localization using functional magnetic resonance imaging (fMRI). Since visual perception unfolds in milliseconds (Thorpe et al., 1996), it is important to elucidate to what extent cortical object and scene processing can be differentiated over the course of neural processing, in addition to cortical regions.

Second, to identify specialized visual processing, researchers typically rely on contrastive experimental paradigms that explicitly separate information across distinct stimulus conditions - for example, contrasting pictures of isolated object cut-outs to those of empty rooms (Epstein and Kanwisher, 1998; Grill-Spector et al., 2001; Harel et al., 2016; Silson et al., 2016). In real life, humans rarely encounter isolated objects or stripped-down scenes; objects are embedded within scenes, and scenes usually contain multiple objects. Importantly, the brain is sensitive to these co-occurrences of objects and scenes in the real-world: guidance of eye movements to congruent and expected objects, or improved scene categorization under difficult conditions, is facilitated by visual cortex co-representing object and scene information (Mack and Eckstein, 2011; Stansbury et al., 2013; Sadeghi et al., 2015; Brandman and Peelen, 2017, 2019; Bonner and Epstein, 2021). This raises the question to what extent specialized processing as identified under contrastive experimental designs is also present during natural visual experience.

An alternative to contrastive designs is to present observers with real-world natural images - containing both background scene information and (multiple) objects - and then identify functional specialization by disentangling how the brain encodes distinct aspects of these images. In fMRI, voxel-wise encoding models (Huth et al., 2012; Stansbury et al., 2013; St-Yves and Naselaris, 2018) have revealed both coarse-grained and fine-grained specialization during natural image perception, allowing identification of features that drive cortical activity in both scene- and object-specific brain regions (Lescroart and Gallant, 2019; Henderson et al., 2023; Adeli et al., 2026). In M/EEG, this approach is comparatively less explored. Early work used linear regression models to study neural encoding of natural scene statistics (Groen et al., 2013; Harel et al., 2016), and recent studies have leveraged encoding models to study visual feature processing during image or short video perception (Cichy et al., 2016; Gifford et al., 2022; Karapetian et al., 2026; McMahon et al., 2026). However, to our knowledge, encoding models have not yet been used to explicitly disambiguate object- and scene processing in temporally-resolved signals.

Representational similarity analysis (RSA) has been used to identify temporal signatures of visual processing during natural image perception, such as low-level spatial features (e.g. GIST) vs. semantic features (e.g., object labels or affordance labels) (Greene and Hansen, 2018, 2020; Dwivedi et al., 2024; Bartnik et al., 2025a). However, since low-, mid- and high-level features of natural images are inherently correlated (Malcolm et al., 2016) and likely build upon each other to aid natural image understanding (Groen et al., 2017), it remains challenging to isolate specialized processing in neural responses to naturalistic real-world natural images (Lescroart et al., 2015). Optimized stimulus sampling (Groen et al., 2018) or statistical decorrelation of hypothesized representational spaces (Greene and Hansen, 2020) helps capture distinct neural variance with RSA, but since RSA relies on relating overall between-image dissimilarity in neural responses to overall model feature dissimilarity, it is less suitable for studying how individual natural image elements (e.g. objects vs. backgrounds) are encoded in neural activity at the single image level. Here, we present an alternative approach: separating objects from scene parts through image-grounded encoding models.

The basis of our encoding models are pretrained image-computable convolutional deep neural networks (DCNNs). In the past decade, DCNNs have emerged as powerful predictors of neural signals across a wide range of measurement modalities (e.g., Schrimpf et al., 2018; Güçlü and Van Gerven, 2015; Gifford et al., 2022). This is because DCNNs learn to extract spatially localized features at multiple levels of complexity, vastly outperforming traditional image-computable models that model one feature space at a time (Yamins et al., 2014; Khaligh-Razavi and Kriegeskorte, 2014; Doerig et al., 2023). Beyond varying the pretrained neural network model to identify the most predictive model architecture (Schrimpf et al., 2018; Conwell et al., 2024; Sartzetaki et al., 2024, 2025), layer (Güçlü and Van Gerven, 2015), or training data (Loke et al., 2025), DCNN-based encoding models further allow explicit manipulation of their visual inputs. For example, we recently used DCNN-based encoding to separate foveal from peripheral information processing in EEG signals (Müller et al., 2026). Others have separated face from scene information to identify functional specialization in visual cortex using fMRI (Ratan Murty et al., 2021).

In the current study, we leverage DCNN-based encoding to explicitly separate *object-bound* from *scene-bound* features in complex natural images. Specifically, we predict EEG responses to high-resolution, large-field intact images (containing both scene background and multiple objects) using distinct encoding models that receive as input spatially masked versions of the images, containing either only pixels labeled as objects, or only pixels labeled as scene background. By comparing the time courses of encoding performance for these two distinct encoding models trained on the same responses, we identify time points during which EEG responses primarily reflect scene vs. object processing. By utilizing activations from all convolutional layers (2D feature maps) of the mutually exclusively-masked DCNNs, we extract potentially relevant features bound to either objects of scenes at multiple levels of feature complexity. Thus, our aim is not to differentiate the level of feature complexity driving neural response selectivity; rather, we focus on the spatial separation of objects vs. scene elements within the image to test for evidence that object- vs. scene-bound features are processed at distinct temporal stages in EEG.

To preface our findings, we observe that scene-bound DCNN features are consistently encoded earlier in the EEG signal compared to object-bound DCNN features. Control analyses confirm that consistent masking of only object or scene pixels is necessary to observe these temporally distinct encoding profiles, and that known center-periphery biased sampling in the EEG signal (Müller et al., 2026) does not explain away the delayed object encoding profile. In follow-up analyses, we use state-of-the-art image segmentation models to go beyond the binary object-scene distinction and identify how features derived from individual object classes and scene elements are encoded in EEG. Specifically, we derive temporal templates of object- and scene-specific visual processing and use these to characterize the typicality of the processing of objects or scene elements in temporally-resolved brain signals.

## Materials and Methods

### Stimuli

To study the neural representations of objects and scene elements embedded in natural scenes, we used 4680 ultra-high resolution photographs from the Open Amsterdam Data Set (OADS) (Müller et al., 2026), depicting Amsterdam street scenes with many objects as they naturally occur in real-world environments. Photographs were taken with an original resolution of 5468×3672 pixels and stored in a uncompressed format and were downsampled to 2155×1440 pixels for this study. Stimuli contained images of streets, buildings, parks, roads with and without cars, bicycles, pedestrians, and other, commonly found, objects of urban environments (**Fig. 1A**). The dataset is publicly available on OSF (see Data availability statement).

**Figure 1:**
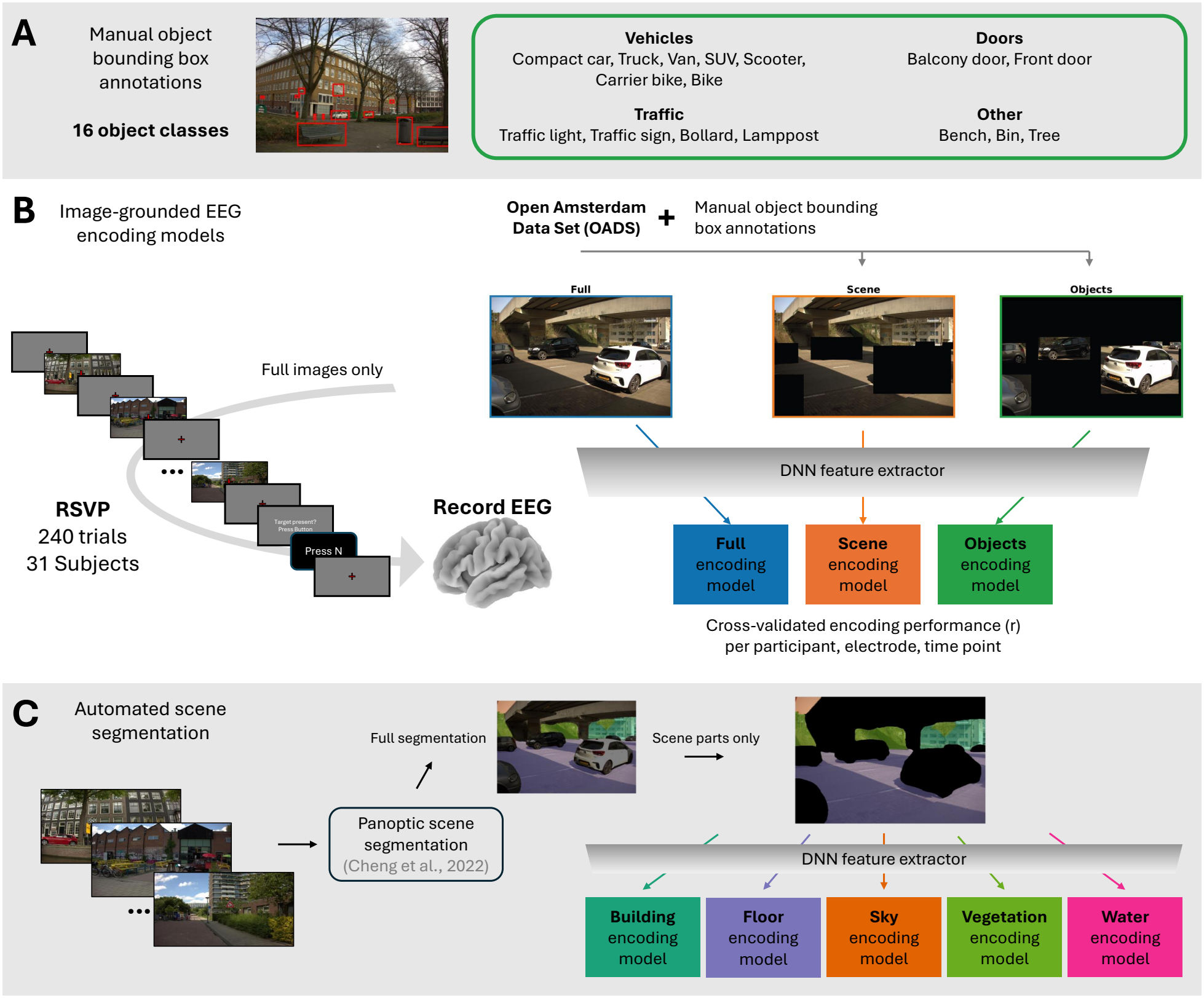
Building image-grounded EEG encoding models for real-world images. **A)** Natural scene images from the Open Amsterdam Data Set (OADS) (Müller et al., 2026) were manually annotated by human observers with bounding boxes for 16 object classes (green box). **B)** EEG data was collected using a rapid serial visual presentation (RSVP) paradigm during which 31 participants were rapidly presented with high-resolution, large-field, real-world street scenes while performing an oddball, indoor-image detection task. We built EEG-predictive encoding models based on DCNN-features for three separate sets of stimuli: original stimuli (“Full”), stimuli from which annotated objects were removed by masking pixels within object bounding boxes (“Scene”), and stimuli where pixels outside of the bounding boxes were masked (“Objects”). **C)** Automatically created image segmentation annotations (Cheng et al., 2022) were used to build EEG encoding models receiving only specific scene elements such as sky, building, or floor information.

**Figure 2:**
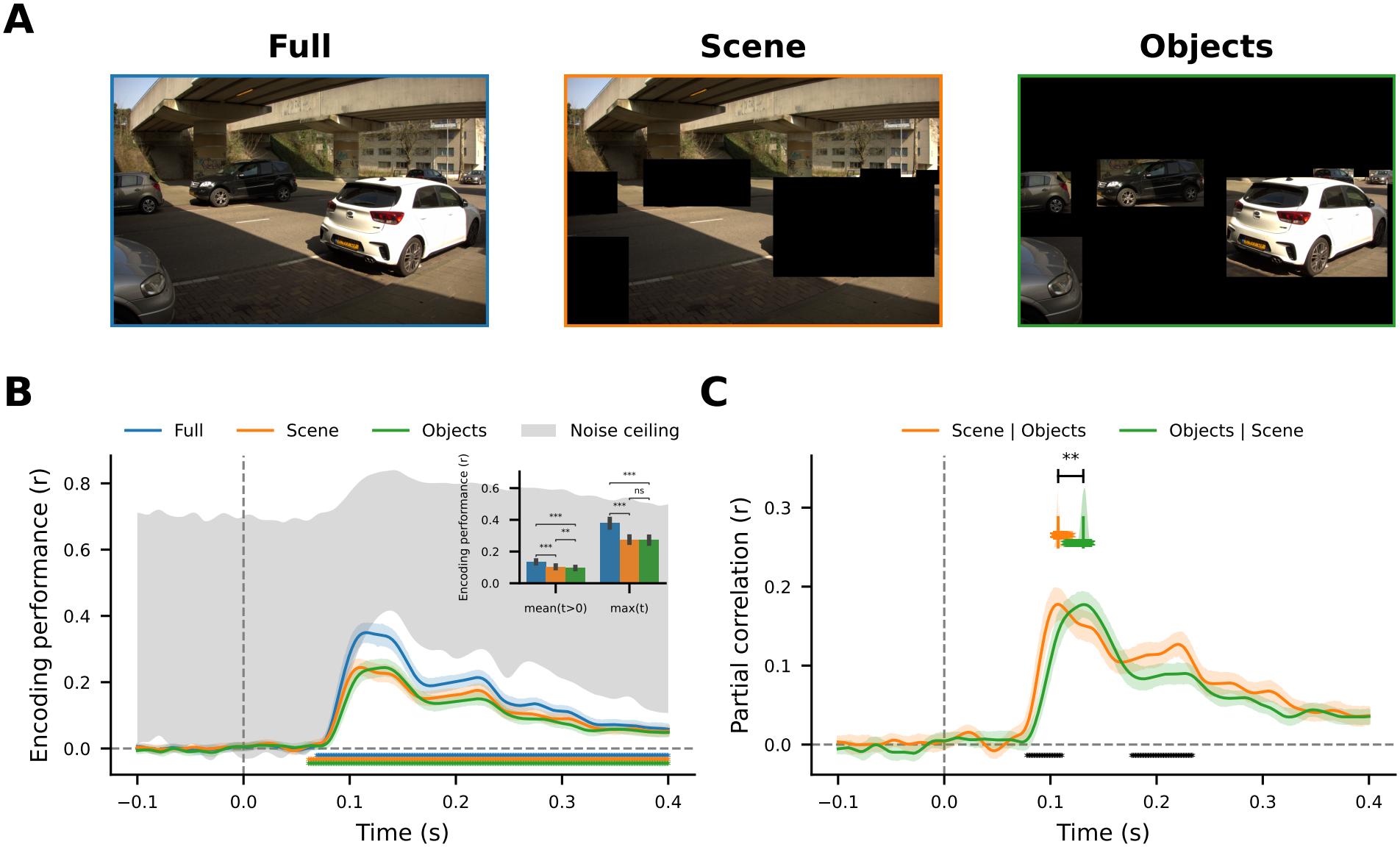
Distinct temporal encoding profiles for scene vs. object information in intact natural image perception. **A)** Example images for “Full image” version, “Scene” version where annotated objects are masked, and “Objects” version where only annotated objects are kept and the remaining pixels are masked, from left to right. **B)** Average encoding performance (correlation) over time for separate models using full, object-only, and scene-only image information, fitted to EEG responses to the intact stimuli. Encoding scores were averaged across all 18 posterior electrodes and 31 participants. Coloured asterisks indicate significant encoding performances across participants, per model, per time point (non-parametric cluster-level t-test, alpha=0.05). Shaded gray areas show the upper and lower bound of the estimated noise ceiling computed across stimulus repetitions on the test set per participant. Inset shows the time-averaged (t>0) and maximum encoding performance per model and asterisks indicate significant differences between models across participants (Wilcoxon signed-rank test, *** p<0.001, ** p<0.01, * p<0.05). **C)** Average partial correlation of predictions of the Scene encoding model with observed EEG responses while regressing out variance explained by the predictions of the Object model, and vice versa. Coloured asterisks and distributions show the time points of maximum partial correlation per bootstrap iteration, vertical bars indicate the means across iterations and black asterisks above indicate statistical differences (p=0.0073; see Methods Statistical analyses). Black asterisks below the lines indicate significant differences per time point between the partial correlations (non-parametric cluster-level paired t-test, alpha=0.05). Encoding results for using multiple different DCNN (varying architecture, training dataset, and training regime) can be found in **Supplementary Fig. S4**.

### EEG experiment

#### Experimental design

To study the temporal dynamics of visual processing of the OADS natural scene stimuli, we used the electroencephalography (EEG) recordings from Müller et al. (2026). In that study, 31 participants were presented with a total of 4680 unique stimuli (702 per participant) in a rapid serial visual presentation (RSVP) paradigm (Grootswagers et al., 2019; Gifford et al., 2022) (**Fig. 1B**, left). Each participant was presented with 240 RSVP trials, in which 20 images were presented for 100 ms each in rapid succession, interleaved with 300 ms blank gray screens. Participants performed a target detection task on interspersed indoor scene images (max. 1 indoor image per trial; half of all trials were target trials) and were asked to respond at the end of each trial via a button press. Target scenes were selected from a separate, custom dataset and were, together with the corresponding EEG responses, excluded from analysis. Stimuli were split into a training (66%, N=468) and a test set (33%, N=234). Following standard approaches for encoding modeling (Naselaris et al., 2011; Gifford et al., 2022), test images were repeated more often than training images to improve their SNR; specifically, training images were repeated 5 times during the EEG experiment and test stimuli were repeated 10 times. Stimuli were presented on a 24-inch monitor with a resolution of 2560×1440 pixels and a refresh rate of 144 Hz. Subjects were seated 63.5 cm from the monitor and were using a head rest such that stimuli spanned 50×29.5 degrees of visual angle. More details about the experimental design can be found in Müller et al. (2026).

#### EEG analysis

EEG data was collected using a Biosemi 64-channel ActiveTwo EEG system (Biosemi Instrumentation) with an extended 10–20 layout. EEG recording was followed by offline re-referencing to external electrodes placed on the earlobes. Resulting EEG data was pre-processed according to our lab’s standard pipeline (Groen et al., 2012) using the MNE software package (Gramfort et al., 2013), and involved filtering (high-pass filter at 0.1 Hz and a low-pass filter at 30 Hz), artifact rejection (removal of deflections *>* 300 *µV*), epoch segmentation in -100 to 400 ms from stimulus onset, ocular correction using electro-oculograms (EOG) electrodes, baseline correction between -100 and 0 ms, and a current source density transformation. Specific details can be found in Müller et al. (2026). The resulting event-related-potential (ERP) data were averaged across repetitions per unique image, thus resulting in an ERP specific to each subject, electrode and image.

#### Noise ceiling calculation

We define the noise ceiling as the maximum explainable variance in the ERP response amplitudes to the test stimuli, computed using a method previously described by Gifford et al. (2022). Essentially, the noise ceiling indicates the reliability of responses across repetitions of the same stimulus. To compute the noise ceiling, we first split the ERP responses for the ten repetitions of the test set stimuli into two non-overlapping splits of 5 repetitions each. We then estimate the lower bound of the noise ceiling as the correlation between the average of the first split of 5 repetitions and the average of the second split of 5 repetitions (split-half reliability), and the upper bound as the correlation between the average of the first split of 5 repetitions and the average of all 10 repetitions. Noise ceilings were computed separately for each subject, electrode and time point.

### Image annotations

#### Manual object annotations

Each OADS image was manually annotated using the free version of the Supervisely annotation tool (https://supervisely.com). Images were downsampled to 1374×918 pixels, converted to JPEG, and then uploaded to the web-based annotation tool. Multiple annotators (the authors and 9 undergraduate students from their lab) first picked a set of object classes by skimming through images and identifying objects with multiple different exemplars and frequent occurrence (see **Fig. 1A** right, for a list of object classes). Then, the annotators were instructed to identify all instances of each object class within each image and to draw the smallest rectangles that included all pixels belonging to the object (**Fig. 1A** left). Areas that were cluttered with multiple overlapping objects were annotated with the special class “MASK” and these annotations were excluded from all analysis (i.e. these areas do not belong to the object image version but are instead included in the scene version). For an overview of the spatial distribution per object class see **Supplementary Fig. S1**.

For each image, we used the manually created annotations to construct multiple stimulus versions that only show information of a certain type. Specifically, we created an Objects version, in which all pixels falling within any bounding box were kept intact, while all other pixels (falling outside of bounding boxes) were masked (set to 0 for all color channels). Additionally, we created a Scene version, in which all pixels falling within any bounding box were masked (set to 0), while all other pixels were kept intact. Last, we also included a stimulus version that uses the original, full, intact image.

#### Automated image annotations

The manually created annotations are focused on object information and were used to globally distinguish between object and scene (i.e., non-object) information and between different kinds of objects. However, these annotations do not differentiate scene elements such as buildings, ground, or sky. While objects often have a largely convex shape and can thus be annotated using simple bounding boxes, scene elements such as buildings or the ground often have complex shapes and are difficult to capture using bounding boxes. Further, alternative annotations such as binary masks or complex polygons are laborious for human annotators. We thus opted to use a state-of-the-art scene segmentation model by Cheng et al. (2022) to obtain pixel-wise labels, covering 71 object and scene part classes (**Fig. 1C**). For each scene element X, we created an additional stimulus version, in which only pixels annotated as X were kept intact while all other pixels were masked. For an overview of the spatial distribution of scene elements see **Supplementary Fig. S2**.

### Computational modeling

To study how different image information is encoded in human EEG recordings, we built linearized encoding models based on the features of a pretrained deep convolutional neural network (DCNN) to predict stimulusevoked event-related potentials (ERPs) at each time point and electrode for each participant. For each of the stimulus versions (full image, objects-only, and scene-only, as well as the additional stimulus versions created for the control analysis and the analyses on individual object classes and scene elements described in the below sections), we built separate encoding models that each received stimuli in which only a specific subset of pixels was kept intact while the remaining pixels were masked (**Fig. 1B**).

#### DCNN feature extraction

We separately extracted the features from a task-optimized AlexNet (Krizhevsky et al., 2012) trained on object classification on ImageNet (Russakovsky et al., 2015) for each stimulus version, i.e., for the full images, the objects-only version (with masked scene-pixels), and the scene-only version (with masked object-pixels). Specifically, we extracted the output of all three max-pool layers (“features.2”, “features.5”, and “features.12”) and used them to build separate encoding models to predict the same EEG data from participants viewing the full, intact stimuli. The resulting encoding performance for each stimulus version was subsequently averaged across layers. Layer-wise encoding performances can be found in **Supplementary Fig. S3**. To test if the choice of DCNN architecture and training regime matters for the outcomes of the encoding model performances we also trained encoding models on the features from an AlexNet trained on scene-categorization on the Places365 dataset (Zhou et al., 2017), an AlexNet trained using a self-supervised contrastive training regime on ImageNet (Zbontar et al., 2021; Prince et al., 2024), a ResNet18, ResNet50 (He et al., 2016), and a VGG16 (Simonyan and Zisserman, 2014) all trained on object classification on ImageNet for which the encoding performances can be found in **Supplementary Fig. S4**.

#### Linearized encoding model

To construct a single encoding model, we performed dimensionality reduction on the extracted DCNN features and fitted a linear regression model, following standard approaches in EEG encoding (e.g., Gifford et al., 2022). We first flattened the extracted features for each stimulus and then applied principal component analysis (PCA), reducing the number of dimensions to 100. Next, we fitted a linear regression model per participant, time point, and electrode, linearly mapping the PC-projected features of the training set stimuli onto the measured, raw ERP amplitudes to those stimuli. We evaluated the model’s cross-validated encoding performance by applying the fitted model to the PC-projected features of the test stimuli and calculating the Pearson correlation (r) between the measured and the predicted ERP amplitudes on those stimuli. The cross-validated encoding performance was averaged across the 18 posterior electrodes (O1, O2, Oz, Iz, P1, P2, P3, P4, P5, P6, P7, P8, PO3, PO7, POz, PO4, PO8, Pz) and participants, and then used to compare different models across time points of visual processing.

#### Encoding model comparison

We use capitalization to refer to the different encoding models, such as “Objects model” for the encoding model receiving the natural scene stimuli with object bounding boxes left intact and background pixels masked out, “Scene model” for the model receiving stimuli with object bounding boxes masked and the background left intact. We compare models in two ways: based on their raw cross-validated encoding performance (r), or based on partial correlations. To compute partial correlation between a model *X* and the measured EEG responses while regressing our the variance explained by a range of other models *Y*_*i*_, we first obtain their predicted responses *P*_*X*_ and 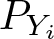, and then compute the correlation between *P*_*X*_ and the measured ERP data while partialling out the variance that is explained by 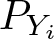.

### Control analyses

We here separate object from scene information within individual images by masking all pixels that lie either inside (object information) or outside (scene information) the object bounding boxes. As a consequence, the used stimuli contain large masked areas with 0 pixel values. To assess the extent to which masking pixels reduces DCNN predictivity of EEG data overall (unrelated to the distinction between object and scene information), we performed several control analyses in which we similarly masked large numbers of pixels, but without explicitly dividing object and scene information.

#### Shifted bounding boxes

We systematically shifted the position of each individual bounding box away from its original location by 20, 40, 60 or 80% until there was no more overlap between the object label and the information the bounding box includes (100%). For each bounding box, the shifting direction was randomly chosen from left or right and up or down. By simply changing the position of each bounding box we maintained the average number of pixels that were masked for a given image (if a bounding boxes shifted outside of the image boundaries, we made it re-appear on the opposite side). For each of these stimulus versions, we then extracted the DCNN features, and fitted new encoding models for either the within bounding box features (“Objects”) and the outside bounding box features (“Scene”) (**Fig. 3**).

**Figure 3:**
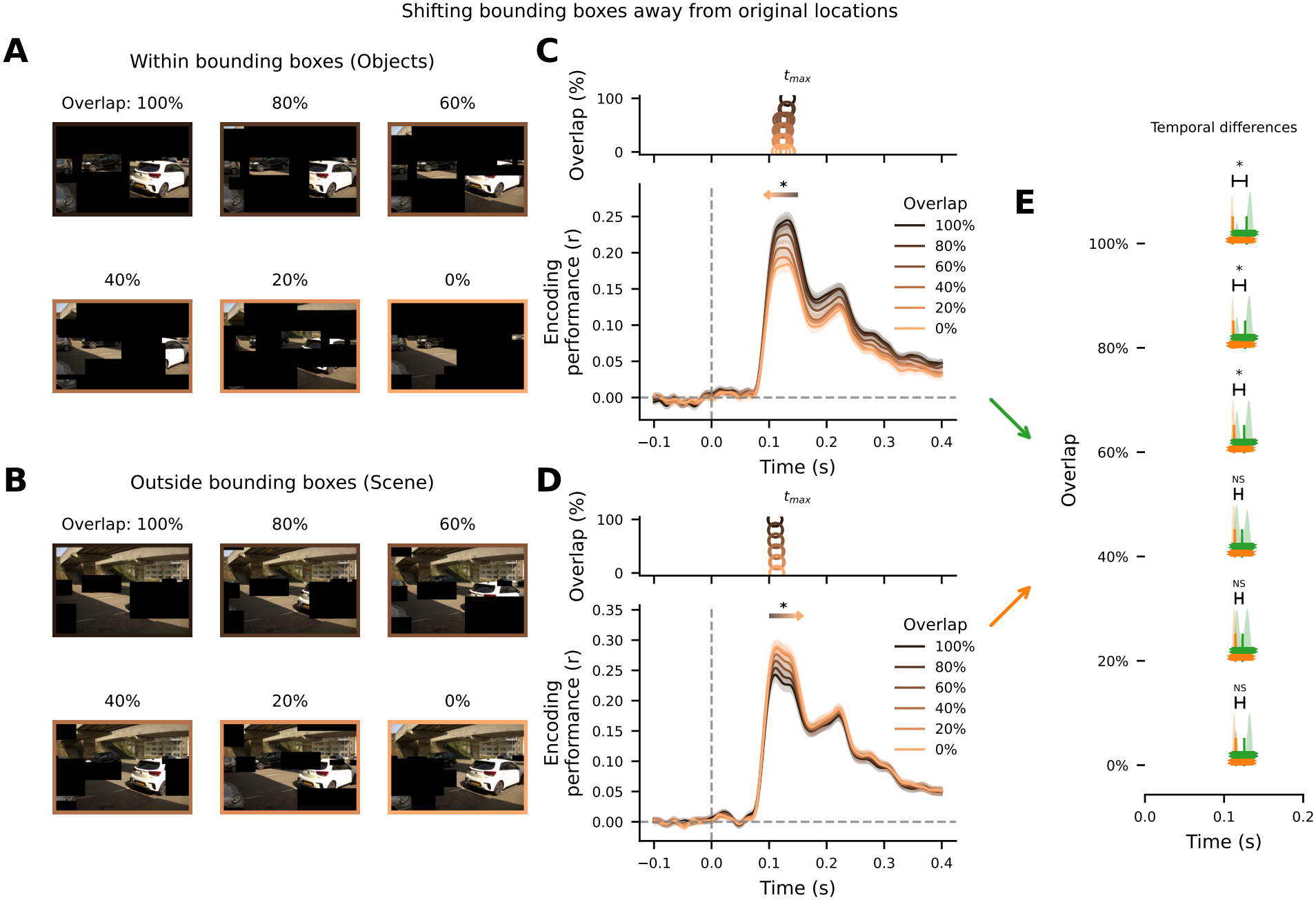
Temporal difference depends on the separation of object from scene information. **A)** Example stimuli for masking pixels outside of bounding boxes after shifting spatially bounding boxes creating stepwise less overlap with their original locations thereby leading to a less distinct separation of object from scene information (100-0% overlap in steps of 20%). **B)** Same as in A, but for the stimulus versions in which pixels within the shifted bounding boxes were masked. **C)** Line plots show the average encoding performance across participants over time for the Objects model version when reducing the amount of overlap of bounding boxes with their original locations. For each participant, the encoding performance was averaged across 5 separate runs with different random seeds as bounding boxes by chance might be shifted to unrepresentative positions (e.g. all overlapping with each other). Colored circles at the top show the average time point of the maximum encoding performance across participants for each of the 5 runs. The colored arrow and black asterisk on top of the line plot indicates a statistically significant shift of the mean time point of the maximum encoding performance in order of reducing overlap. **D)** Same as in C, but for the Scene model version. **E)** Distribution of the timepoints of maximum encoding performance for each model shown in C and D, vertically ordered by the amount of overlap of shifted bounding boxes with their original locations. Coloured asterisks and distributions show the time points of maximum encoding performance per bootstrap iteration, vertical bars indicate the means across iterations and black asterisks on the indicate statistical differences of each pairwise comparison between the Scene and Objects model versions.

**Figure 4:**
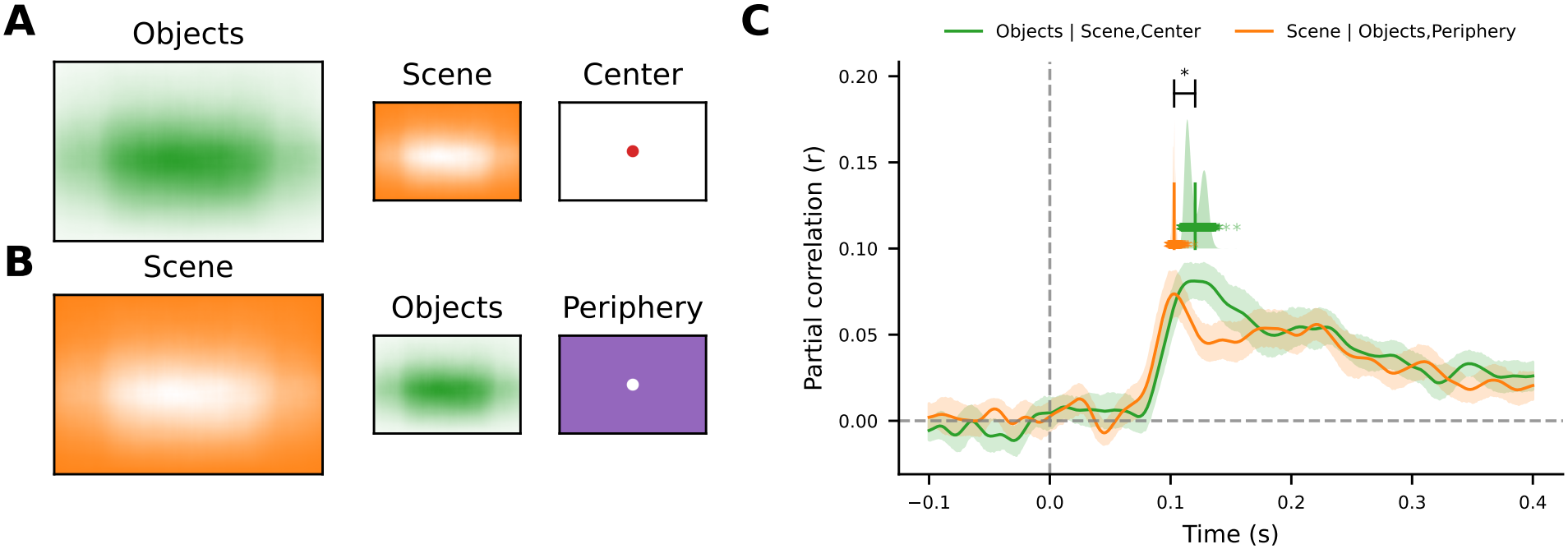
Object encoding is distinct from foveal processing. **A)** Illustration of the summed spatial distribution across all object annotations and all images (left), all scene information (middle) and the image area kept intact for the Center model (right). For a detailed quantification of the per-class spatial distribution see **Supplementary Fig. S1. B)** Same as in A), but for the distribution of scene information (left), object annotations (middle), and the image area kept intact for the Periphery model (right). **C)** Line plots show the average partial correlation over time across participants for the Objects model while regressing out the variance explained by the Scene model and a model receiving only information from central and thus foveal regions of the image (green line) and for the Scene model while regressing out the variance explained by the Objects model and a model receiving only information from the periphery of the image (orange line). Coloured asterisks and distributions show the time points of maximum partial correlation per bootstrap iteration, vertical bars indicate the means across iterations and black asterisks above indicate statistical differences (p=0.0073; see Methods). Black asterisks below the lines indicate significant differences per time point between the partial correlations (non-parametric cluster-level paired t-test, alpha=0.05).

#### Matching the number of scene and object pixels

In our stimulus set, the number of pixels annotated as objects is smaller than the number of remaining scene pixels, creating an imbalance in the number of intact pixels in the stimuli that are used as inputs to the Objects encoding model and the Scene encoding model. To test if this difference in number of pixels could be a reason for observing differences between the Objects and Scene encoding performance, we additionally tested versions of the Scene model, in which we systematically reduced the number of scene pixels that were kept intact and then tested the resulting encoding performance against that of the Objects model (**Supplementary Fig. S5**).

#### Applying bounding boxes on DCNN features instead of images

By applying the bounding boxes on the images that are fed to the DCNN to extract its features, large black areas are created in the DCNN input images that potentially change the way the CNN processes the Objects version and the Scene version of the same image. To ensure that the potential differences of this are not the sole reason for any differences in encoding performance between the Objects and the Scene model, we trained additional encoding models in which we first fed the intact, full image to the CNN, extracted its features and only then applied the bounding boxes to the extracted 2D DCNN feature maps to create the Objects and the Scene versions (**Supplementary Fig. S6**).

#### Randomly assigned bounding boxes

To test if the overall spatial distribution of the bounding boxes across images is the key reason for any observed differences between the Objects and the Scene version, we included a version in which we randomly assigned bounding boxes to other images, such that the location of bounding boxes does not meaningfully map onto the location of the actual objects within images. If the encoding performances are similar to the original performance that used true bounding boxes, then it might not be the content of the individual bounding boxes per image that is important for observing differences between the Objects and Scene version, but rather the overall distribution of bounding boxes (**Supplementary Fig. S7**)

#### Foveal and peripheral information

In our stimulus set, most annotated objects are positioned around the horizontal meridian (**Supplementary Fig. S1**). By design of the EEG experiment, the center of the images is also the point of fixation of the participants. Thus, information at the center of the images falls within foveal regions of the retina which, as we showed previously (Müller et al., 2026), dominates the EEG signal at posterior electrodes. To test whether this overlap of object information with foveal regions could explain the observed temporal differences in encoding profiles for the Object and Scene models, we included additional encoding models using stimulus versions in which either only foveal information was maintained (“Center”), while peripheral information was masked, or vice-versa (“Periphery”). Similarly to separating object from scene information, we separated foveal from peripheral information by either masking all pixels at the center of the image (1% of the total image area) or by keeping those pixels intact and masking all other pixels. We then extracted the DCNN features from either center or periphery and fitted linear encoding models to obtain model predictions for stimuli in the held out test set. We then quantified the amount of variance explained by object information, independent of both foveal and scene information, by computing the partial correlation of the Objects model predictions with the measured ERP responses while using the Center model predictions and the Scene model predictions as covariates. Similarly, we quantified the variance explained by scene information, independent of peripheral and object information, by computing the partial correlation of Scene model predictions with ERP responses while using the Periphery model predictions and Objects model predictions as covariates.

### Individual object classes and scene elements

#### Encoding contribution of object classes and scene elements

To investigate the contribution of individual object classes to the EEG predictivity, we included two additional encoding models per object class. For each object class X, the first encoding model received stimuli showing only objects except for objects of class X (those pixels were masked as well). The second encoding model received stimuli showing all scene information plus objects of class X, while pixels belonging to any other object class were masked. We denote these image versions as “Objects - X” and “Scene + X”, where X is one of the 16 object classes. Additionally, for each object class, we summed the number of pixels across images belonging to that object class and divided them by the total number of object pixels across all classes. We then related the change in EEG predictivity when including or excluding an object class to the proportion of pixels that that class takes up across all images (**Fig. 5**).

**Figure 5:**
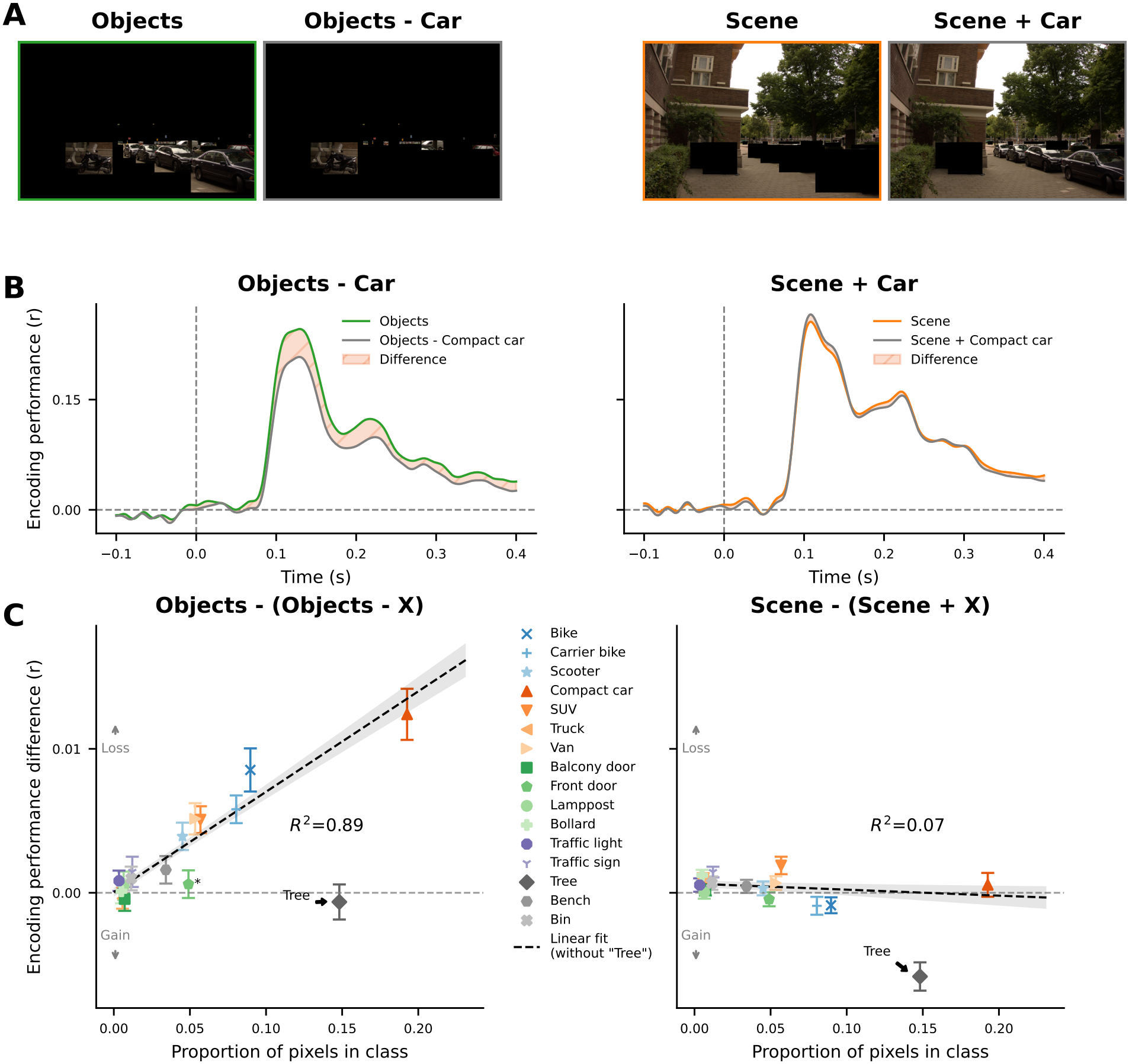
Linear relation between number of pixels and encoding contribution of individual object classes - except for Trees. **A)** Example stimuli for the Objects model (showing all objects), the Objects model without cars, the Scene model (showing everything that is not an object), and the Scene model plus cars, from left to right. **B)** Illustrative plots of the encoding performance (r) for the Objects model and the Objects model without cars (left plot) as well as the Scene model and the Scene model plus cars (right plot). Shaded areas show the difference between the encoding performances of each pair of models. **C)** The difference in encoding performance (r) as illustrated in B), for each individual object class plotted against the proportion of pixels that the object class takes up across all images. Left panel shows the difference in encoding performance between the Objects model and the Objects model without each individual object class and right panel shows the difference between the Scene model and the Scene model with each individual object classes added. Error bars show the standard error per object class across participants. Dashed lines indicate the best linear fit of all classes except the class “Tree”, which forms an outlier, suggesting it is not encoded as an object in the EEG signal. Gray shaded areas indicate the standard error of the linear fit across participants. No classes were identified as outliers from the linear relationship after excluding “Tree” (see Methods).

Similarly, for each scene element X identified by the automated image segmentation (Cheng et al., 2022), we compared the encoding performance of a model receiving stimulus showing all scene information (with masked object information) except for pixels of the element X (those pixels were masked as well). We then related the change in encoding performance to the proportion of pixels that that elements takes up across all images (**Fig. 6**).

**Figure 6:**
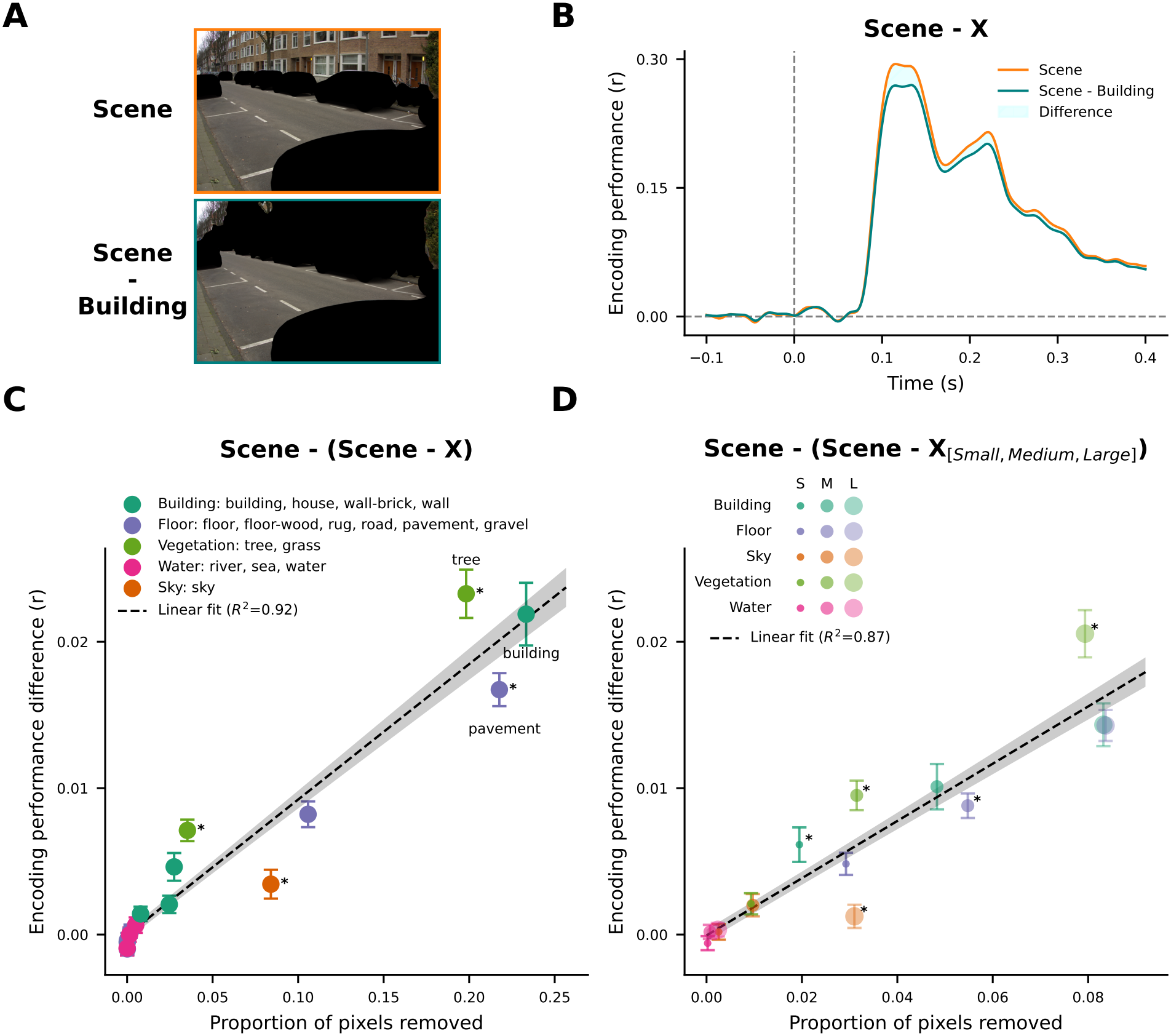
Contribution of scene elements is largely defined by their spatial extent. **A)** Example image with all object information masked (top) and additionally one scene element (building) masked (bottom). **B)** Illustration of how we identify the encoding profile of a single scene element: we compare the encoding performance of the original Scene model to that of a model receiving images where pixels belonging to a specific scene element were masked. The encoding profile of that scene element is then quantified as the temporally-averaged difference between the two encoding performances (shaded area). **C)** Difference in encoding performance (r) as computed in B) for each individual scene element relative to the proportion of pixels that the scene element takes up across all images. Error bars show the standard error per scene element across participants. Dashed lines indicate the best linear fit across all scene elements. Gray shaded areas indicate the standard error of the linear fit across participants. Asterisks indicate scene elements identified as outliers from the linear relationship (see Methods). **D)** Same as in C, but for repeating the analysis with specific sizes of each scene element.

#### Scene part sizes

To study how the size of annotated scene elements links to the EEG encoding performance, we divided the stimuli for each of the scene elements into subsets of images containing either large areas of that scene element, i.e. many pixels, or medium-sized areas, or small areas, i.e. few pixels. Sets were divided by first ordering stimuli according to the number of pixels per scene element and then splitting them into the top, middle, and last percentile (**Fig. 6D**).

### Statistical analyses

For all analyses comparing time point-wise significant difference of encoding performances (correlations or partial correlations) (between models or against 0; **Fig. 2B, C and Fig. 4C, D**), we performed non-parametric cluster-level permutation t-tests across participants for each time point using alpha=0.05 (Maris and Oostenveld, 2007; Sassenhagen and Draschkow, 2019). For the analysis comparing time-averaged encoding performances between models (inset in **Figs. 2B**), we performed Wilcoxon signed-rank tests across participants with FDR-correction (Benjamini and Hochberg, 1995).

To compare whether encoding models show differences in the time point of maximum encoding performance, we used bootstrapping across participants, for robust estimations of peak encoding performances. Specifically, across 10000 bootstrap iterations, we randomly sampled with replacement from the set of 31 participants, then averaged the encoding performances within the sample and identified the time point of the maximum, for each encoding model. More specifically, we took the first time point after stimulus onset at which the encoding performance is both a local maximum and at least half of the global maximum. We then compared the variance of the peak time points across encoding models against the variance of a null distribution by first shuffling the order of encoding models before computing the variance, yielding a p-value (number of iterations with equal or larger variance of the null distribution compared to the observed variance). We used this bootstrapping method to compare the peak time points of the partial correlations between the Object and Scene models in **Figs. 2C, 4C**, and **Supplementary Figs. S3A, S4, S6D**, and the raw encoding performances of models in **Fig. 3E**, and **Supplementary Figs. S5C, S7C, S8B**. Additionally, to test for a monotonic trend of the timepoint of the peak encoding performance across the parameterized conditions of the bounding box shift analysis (**Fig. 3C-D**), we used Page’s L trend test on the peak encoding performances across subjects and across shifting conditions, assuming the following orders: [0, 20, 40, 60, 80, 100] % shifted for the Objects model conditions and [100, 80, 60, 40, 20, 0] % shifted for the Scene model conditions.

To test whether object size is predictive of the class’ contribution to the encoding performance, we first compared the encoding performance of a version of the Objects models where additionally the pixels of one object class were masked to that of the Objects model (**Fig. 5B**). We then related the relative differences in encoding performance for each individual object class to the proportion of pixels that each class takes up. Similarly, we tested how the encoding performance changes between the Scene model and models that receive all scene information plus the pixels from one object class (**Fig. 5C**) as well we how removing individual scene elements obtained using automated image segmentation changes the encoding performance compared to using all scene information (**Fig. 6C-D**). For all three analyses, we quantified the relationship between proportion of masked pixels and relative change in encoding performance using a linear regression per participant. To test whether any object classes or scene elements form an outlier in this linear relationship, we performed a Wilcoxon signed-rank test for each object class and scene element, testing if the residuals across participants are significantly larger than 0 (alpha=0.05; fdr-corrected across all object classes or all scene elements; outliers are highlighted with asterisks in the figures).

## Results

The goal of the current study was to identify temporal signatures of scene and object processing in the human brain during naturalistic vision. To this end, we developed image-grounded encoding models which we applied to a large-scale dataset consisting of human EEG responses to high-resolution, natural images of real-world, urban environments. First, we manually created bounding box annotations for a set of 16 object classes, identifying for each image which objects are present, where they are located, and their spatial extent (**Fig. 1A**). Second, for each image, we divided pixels into two groups - objects or scene, whereby all pixels contained within the bounding boxes were defined as objects and all remaining pixels were defined as scene. Then, we used these two groups of pixels to create new stimulus versions that showed either only the object pixels, or only the scene pixels, while all other pixels were masked (set to 0; **Fig. 2A**). For each stimulus version, we extracted the convolutional activation maps of a pre-trained DCNN (the three maxpool layers of AlexNet; see Methods DCNN feature extraction). For the stimuli with masked scene pixels, this resulted in extraction of *object-bound* DCNN features, while for stimuli with masked object pixels, it resulted in extraction of *scene-bound* DCNN features. Using these separate activation maps, we then fitted separate encoding models (referred to as “Objects” and “Scene” models) to predict EEG data recorded from human participants viewing the full, intact stimuli. For comparison, we also built encoding models for the original, unmasked images (“Full”). Encoding models were trained to predict event-related potential (ERP) amplitudes for each participant, electrode, and time point separately, and predicted responses were evaluated by correlating them to the true observed responses on a separate test set (**Fig. 2B**).

### Temporally distinct processing of object and scene information

We first consider raw encoding performance for the Full, Scene and Object models. Cross-validated encoding performance (r) for the Full model rises quickly starting at 80 ms after stimulus onset and peaks at 110 ms, showing that the EEG responses are robustly predicted from DCNN features (**Fig. 2B**). In comparison, the Objects and Scene models have a lower encoding performance. This is expected since for these models, parts of the images are masked, thus part of what participants saw on the screen during the experiment is not included in the encoding model. Time-averaged Scene model performance (left bar plot in inset) is slightly higher than that of the Objects model (Wilcoxon-signed rank test: W(31)=96, p=0.0022). However, peak encoding performance (right bar plot inset) between the Objects and the Scene model is similar, suggesting that both encoding models capture relevant information contained in the EEG responses. Importantly, the temporal dynamics of the encoding curves for the Scene and Objects models diverge; at early time points (∼100 ms), the Scene model appears to explain more variance than the Objects model, while at slightly later time points (∼150 ms) both models seem to perform equally well, followed by a period during which the Scene model has the best performance of the two.

To directly compare the temporal profiles of scene and object encoding, we computed partial correlations between model predictions, removing their shared variance in predicting the EEG recordings (see Methods Encoding model comparison). In this analysis, temporal differences between the Scene and Object encoding model are more clearly pronounced (**Fig. 2C**); scene information explains more unique variance at early time points (cluster-level permutation t-test: significant differences for t=[80-110] ms, p=0.0125) and a later time points (t=[175-235] ms, p=0.0096), while in the intermediate period (110-175 ms), object and scene information perform similarly. Moreover, peak encoding performance is significantly earlier for the Scene model compared to the Objects model (p=0.0114; see Methods Statistical analyses for details of bootstrapping the time point of the peak). These differences are robust across multiple DCNN architectures, training datasets, and training regimes (**Supplementary Fig. S4**).

These results suggest that scene and object information are processed across distinct neural timescales. The early advantage of the Scene model is consistent with prior accounts of precedence of global information in visual information processing (Greene and Oliva, 2009), and reminiscent of a coarse-to-fine processing hierarchy (Navon, 1977; Bar, 2004; Hegdé, 2008), which we here see reflected in the temporal encoding profiles of EEG responses to real-world images. However, before these distinct encoding profiles can be confidently attributed to scene vs. object information processing, several potential alternative explanations need to be addressed.

### Temporal difference depends on the separation of object from scene information

The separation of object from scene information was operationalized as a masking of pixels from which *object-bound* and *scene-bound* DCNN features were extracted. To test whether this masking alone, rather than separating object and scene information, drives the observed temporal differences, we ran several control analyses.

First, we systematically shifted the bounding boxes away from their original location, stepwise reducing the overlap they have with the actual objects. For each shift condition (0-100% in steps of 20%), we ran two models: one that received pixels within the shifted bounding boxes (“Objects”, **Fig. 3A**) and one that received all pixels outside the shifted bounding boxes (“Scene”, **Fig. 3B**). Critically, these ‘shifted’ models receive the same number of pixels as the original Objects and Scene models, allowing us to address whether observed temporal shifts are dependent on true object-vs. scene-bound pixel selection, or can be observed when feeding any sub-selection of pixels to distinct encoding models. We find that for shifted models, timepoints of the peak encoding performance significantly shift with decreasing overlap to the original bounding boxes (**Fig. 3C+D** top). For the Objects model, the peak shifts to earlier timepoints (Page’s L trend test, L=2426.5, p<0.0001), while for the Scene model the peak shifts to later timepoints (L=2777.0, p<0.0001). As a consequence of these shifts in peaks, the temporal difference between the Objects and the Scene models disappears with decreasing overlap with the original bounding box (**Fig. 3E**). This shows that the observed temporal differences do not emerge from separating pixels *per se*, but require separating object-bound vs. scene-bound pixels.

A second potential confound is that the Objects model receives fewer pixels overall compared to the Scene model, potentially leading to an unfair comparison between the two models. The raw encoding performances were indeed slightly higher overall for the Scene model (**Fig. 2B**, left inset), but peak encoding performance was statistically indistinguishable between the Objects and the Scene models, despite the Objects model receiving fewer pixels (**Fig. 2B**, right inset). The bounding box shifting analysis directly addresses this potential confound: shifting bounding boxes away from the original object locations *increases* Scene model performance but it *decreases* Objects model performance (**Fig. 3C,D**), despite the same number of pixels being masked in all conditions. Thus, receiving fewer pixels does not automatically decrease performance: rather, it depends on which specific pixels are included. Consistently, an additional control analysis shows that selecting similar (or even fewer) numbers of scene pixels as object pixels significantly decreases Scene model performance while the temporal difference with the Objects model remains intact (**Supplementary Figure S5**).

Third, masking the input images creates large black regions with sharp, edge-like boundaries that likely affect the features computed by the DCNN. To rule out that the observed temporal differences in object and scene encoding were driven by additional edges that influence DCNN computation, we trained additional encoding models for which the DCNN received the full intact image and masks were applied to the extracted 2D activation feature maps to create the Objects and the Scene version (**Supplementary Fig. S6A-B**). These two models show the same EEG encoding profiles as encoding models trained with masked input images, suggesting that the temporal differences between the Objects and the Scene models were not an artifact of DCNN processing of image masks, but instead signify during what time in the ERP time course object and scene information are encoded (**Supplementary Fig. S6B-C**).

Last, to rule out that the observed temporal differences are a consequence of the overall distribution of e.g. the spatial extent, contiguity or spatial location of the bounding boxes - independent of the content of the bounding boxes - we trained additional encoding models for which the pairing of images and bounding boxes was randomized (see Methods Randomly assigned bounding boxes). In this condition, the exact same bounding boxes are used across all images, yet the bounding boxes do not meaningfully map onto the objects in the images (**Supplementary Fig. S7A**). As a result, pixels within the bounding boxes may depict either objects or scene elements or combinations of both. Thus, we do not expect that the temporal difference between the Objects and Scene model version in this analysis disappears entirely, but that this difference reduces in magnitude. For encoding models trained on randomized image-mask pairings, two patterns change relative to the original Objects and Scene models: first, overall encoding performance of the “Objects” model (i.e., the model receiving information within bounding boxes) reduces drastically, to about half of the original Objects model (**Supplementary Fig. S7B**). Second, the temporal difference between the “Objects” and “Scene” models disappears for the raw encoding performances and is much reduced in partial correlation results (**Supplementary Fig. S7C**). These results suggest that the temporal differences between object and scene processing are not caused by the overall distribution of spatial extent, contiguity or spatial location of the object bounding boxes, but instead depend on the content of the bounding boxes.

Overall, these control analyses indicate that the observed temporal distinction between the Scene and Objects model is not driven by the application of image masks, differences in number of pixels, confounds in DCNN feature computation or distributions of bounding boxes. Rather, they suggest that observing distinct temporal encoding profiles in EEG signals requires clean separation of scene-vs. object-bound features in the input image.

### Distinct encoding of object and foveal information

The analyses above did not yet address a final potential confound related to differential center-periphery processing in the visual system. It is well-known that the retina samples information at the center of gaze more densely (Rodieck, 1979; Oyster et al., 1981) and that this foveal information is overrepresented in early visual cortex (Cowey and Rolls, 1974; Wässle et al., 1989; Kwon and Liu, 2019). In a previous study on the current dataset (Müller et al., 2026), we demonstrated that EEG signals recorded during natural image viewing are indeed also biased by foveal information, with contributions of central and peripheral peaking at distinct time-points in the EEG encoding profile, similar to the scene- and object profiles observed here (peripheral information preceding central information). In our stimulus set, objects were unequally distributed across the scene, reflecting their typical locations in the real-world (**Fig. 4A** and **Supplementary Fig. S1** for heatmaps showing the frequency of appearances per object class for each pixel). Some object classes were overrepresented in the center of the image, which was processed in the fovea, while most scene information was present in the periphery. Thus, it is conceivable that differences in encoding of object-vs. scene-bound features in EEG signals may reflect differential spatial sampling, rather than signatures of functional specialization.

To directly test whether the observed differences in temporal profiles between object and scene information were driven by a systematic difference in fovea-periphery sampling, we calculated the partial correlation of the Objects model with the Scene model but also with an additional encoding model that only received pixels from the center of the image (see **Fig. 4A**; same as “Center” model in Müller et al. (2026), see Methods Foveal and peripheral information). Similarly, we compute the partial correlation between the Scene model and the Objects model and an additional encoding model that only received peripheral image pixels (**Fig. 4B**, “Periphery” model).

**Fig. 4C** shows the time courses of the partial correlation for the Object model, while removing the overlap in predictions with the Scene model and the Center model, and for the Scene model while removing the overlap in predictions with the Objects model and the Periphery model. In this comparison, any temporal difference due to differential sampling of foveal and peripheral information has been accounted for. Nevertheless, we still find that Scene model encoding peaks earlier than Object encoding (p=0.0203, bootstrapping of peak time point). Still, the overall partial correlation reduces to above half of the original Objects and Scene models. These results suggest that while the processing of foveal and peripheral information partially overlap with processing of object and scene information, this overlap is not the sole driver of the observed temporal differences.

### Templates for object and scene processing reveal typicality of object classes

So far, we demonstrated separate temporal encoding profiles for object and scene information within naturalistic, real-world environments. By testing various control models, we verified that these profiles are signatures of the processing of *object-bound* and *scene-bound* features. However, these profiles averaged performance across many object classes. Next, we asked whether it is possible to delineate distinct encoding profiles for the individual object classes contained in our dataset.

For each individual object class X - for example, cars - we built two new encoding models receiving either 1) images containing all objects *except* cars (“Objects - X”), or 2) images containing all scene information *plus* cars (“Scene + X”), but no other objects (**Fig. 5A**). We then compared the resulting encoding performances of these two models to that of the original Objects and Scene models, respectively (**Fig. 5B**). Considering the original Objects and Scene encoding time courses as representative ‘templates’ of overall object- and scene-specific information processing, we reasoned that systematic comparison of the new reduced (1) or enhanced (2) models to these templates could reveal class-specific information encoding, according to the following logic: if an object class was perceived and thus encoded much like any other object, then the removal of this object class should lead to a decrease in Object model encoding performance that is roughly proportional to the area that instances of this class take up in the images (measured as the proportion of pixels of one object class compared to all object classes). At the same time, adding this object class to the Scene model should not lead to a large change in encoding performance. In other words, if a specific object class was processed as a ‘true’ object, its removal from the model should reduce overall Objects model encoding performance, but not affect Scene model performance. However, if an object class is perceived more like a part of the scene, the opposite pattern should be observed.

We find that all but one object class follow the predicted linear relationship between the proportion of pixels and the decrease in encoding performance resulting from removing those objects from the Objects model (**Fig. 5C**, left; R^2^=0.89). Only when removing information labeled as “Tree” did encoding performance not decrease linearly with the proportion of pixels - instead it *increased* Object encoding performance. Additionally, the “Front door” class was identified as an outlier from this linear relationship (Z=75, p<0.0001; see Methods Statistical analyses for how outliers were defined).

Consistently, all but the same one class “Tree” followed a constant relationship between the number of pixels and the change in encoding performance resulting from adding specific object classes to the Scene model (**Fig. 5C**, right; R^2^=0.07). Inverse mirroring the observed increase in Object model performance due to removal of “Tree”, adding “Tree” to the Scene model enhanced its encoding performance. This suggests that while trees (and to a lesser extent, front doors) were here labeled as objects, they are neurally processed as parts of a scene. At a broader level, these results suggest that deviation in image-grounded model encoding performance for a specific individual object class relative to overall Object or Scene encoding can be used as an index of the ‘typicality’ of neural processing of that object class.

### Scene processing template reveals decoupling of spatial extent and neural relevance

In all our analyses so far, we considered individual object classes and defined scene information as all scene parts not contained within an object bounding box. However, just like objects, scene background can be subdivided into separate elements, such as buildings, sky or floor. Next, we investigated whether it is possible to identify individual processing signatures for such elements as well.

Using a pretrained scene segmentation model Cheng et al. (2022), we obtained pixel-wise annotations for each image, now also including segmentations of scene parts (e.g., buildings, sky, or floor) in addition to objects. To verify the quality of annotations, we first selected those object annotation labels that were similar to the ones that were used for our original manual bounding boxes, and compared the encoding performance of a model receiving pixels belonging to the automatically-segmented objects with the encoding performance of a model receiving the pixels belonging to the manually-annotated objects (**Supplementary Fig. S8**). Both models yield similar encoding performances, although automated annotations lead to slightly lower encoding performance. Still, this confirms that the automatically created segmentations were of comparable quality as our manual annotations.

After having established the quality of automatic image segmentations, we built additional encoding models that receive as input all pixels belonging to scene parts, except from pixels belonging to one specific scene part (see example in **Fig. 6A**), analogous to the analysis in **Fig. 5**. Then, we compared the resulting encoding performances of removing the information of a specific scene part to that of a model receiving all scene information (**Fig. 6B**). As before, we related the difference in encoding performance to the proportion of removed pixels.

In general, the difference in encoding performance is proportional to the amount of pixels that were removed (**Fig. 6C**, left; R^2^=0.92). Some scene parts such as Water, contribute very little to the encoding performance but also only occupy few pixels in the images. Other classes significantly deviate from the linear relationship, and either contribute super-linearly (Grass, outlier test: Z=30, p<0.0001; Tree: Z=77, p=0.0005) or sub-linearly (Sky, Z=48, p<0.0001; Pavement, Z=103, p=0.0036). Overall, the strongest contributions to encoding performance appear to be driven by the scene elements Pavement, Tree and Building. The fact that the contribution of some scene elements is disproportional to the number of pixels they occupy may reflect higher neural relevance of these elements, while for others the number of pixels is largely predicting their overall encoding contribution.

To further examine the relation between the number of pixels of an individual scene part and its encoding contribution, we ran the previous encoding analyses at a more fine-grained granularity, selecting for each individual scene part either only all images with many pixels of that scene part (as a proxy for a large area), a medium amount (medium area), or few pixels (small area), which can be seen as a proxy for retinal size. If number of pixels is sufficient to predict the contribution of a scene part to the overall encoding performance, then the different size conditions should reveal a similarly linear relationship between the proportion of pixels removed and the difference in encoding performance. However, if the content of some scene parts is more important to the encoding performance of the Scene model, then the small conditions should have super-linear contributions.

As expected, the change in encoding performance when masking pixels of scene elements of different sizes generally follows a linear relationship with the proportions of removed pixels (**Fig. 6D**; R^2^=0.87). However, there are some interesting deviations from this regularity for specific scene elements. For example, small buildings (Z=117, p<0.009), medium-sized vegetation patches (Z=85, p=0.001), and large vegetation (Z=62, p<0.0001) contribute significantly more to the encoding performance than predicted by a linear relationship, whereas large sky (Z=13, p<0.0001), and medium-sized floor patches (Z=122, p=0.012) contribute significantly less. Overall, these results show that different scene elements, despite occupying similar numbers of pixels, have different encoding contributions, suggesting a decoupling of neural relevance from the spatial extent of individual scene parts.

## Discussion

The human brain is thought to process incoming visual information efficiently through dedicated networks that are specialized for specific types of information, such as objects, faces, or scenes. While the spatial clustering of these networks into topographic regions has been studied extensively using fMRI, the temporal dynamics of specialized information encoding are less well understood. In addition, most prior research has relied on the use of artificial stimuli to separate individual visual components, limiting ecological validity. To identify whether the brain differentially encodes objects and scene elements over time we used a large-scale EEG dataset of temporally-resolved responses to large-field naturalistic images, and built neural encoding models of object and scene information processing. By separating object from scene information at the pixel level and fitting separate encoding models based on corresponding *object-bound* and *scene-bound* features, we identified distinct temporal profiles of object and scene encoding. Additionally, we elucidated how individual components of real-world environments - individual object classes and scene elements - are encoded during visual processing. Our results reveal that when humans perceive complex, realistic natural scenes, scene features are encoded earlier than object features, suggesting their representations involve distinct computational cascades. Our fine-grained analyses furthermore suggest that not all scene elements are ‘perceived equally’: instead, certain image elements are overrepresented compared to others, disproportional to the number of pixels they occupy in the image.

### Temporal profiles of object and scene processing

Previous work has suggested that the P2 ERP component can act as a marker of scene-specific processing (Harel et al., 2016). Here, we instead find that scene encoding peaks at 80-110 ms, preceding peak object encoding. This is consistent with theories of coarse-to-fine processing (global scene context precedes local object classification) (Bar, 2004; Hegdé, 2008), M/EEG studies finding discriminable scene category representations at occipital electrodes as early as 84 ms (Lowe et al., 2018; Ramkumar et al., 2016), and behavioral studies demonstrating ultrarapid recognition of scenes from global image properties (e.g., Greene and Oliva, 2009). The difference in outcomes with Harel et al. (2016) could be due to the use of naturalistic images, i.e. stimuli that contain both scene and object information, compared to their use of isolated stimuli for each condition, as well as the focus of the EEG analyses (ERP components versus encoding of DCNN features). Prior studies have found substantial differences between naturalistic and artificial experimental conditions (Kayser et al., 2004; Hasson et al., 2010; Martin and Schröder, 2013; Talebi and Baker, 2012; David et al., 2004); whether naturalistic stimuli should be preferred over artificial ones when studying the neural mechanisms underlying visual perception continues to be debated (Felsen and Dan, 2005; Rust and Movshon, 2005; Zhang et al., 2021).

More recent work has leveraged representational similarity analysis (RSA) to investigate temporal differences in natural image information processing. Consistent with our results, Greene and Hansen (2020) found that image-computable low-level global scene features around 100 ms preceded encoding of high-level object features around 170 ms in EEG responses. Similarly, using behavioral annotations, Mononen et al. (2025) showed that spatial structure predicts neural signals better at early time points compared to semantic content and vice-versa for later time points. Both these studies relied on whole-image descriptors to identify differences in neural dissimilarity across stimuli (i.e., each feature model compared in RSA is based on the entire image), and thus did not differentiate between separate features within a single scene. In contrast, our image-grounded encoding models disentangle the encoding of scene information and object information within each unique stimulus, allowing for more precise tests of which specific image elements are driving variance in neural responses to natural images.

Prior work has also used DCNNs trained with different task objectives (e.g., object recognition vs. scene classification, Cichy et al., 2017) or with different visual ‘diets’ (e.g., segmented objects vs. objects+backgrounds, Loke et al., 2025) to obtain object- and scene-features from images and search for temporal signatures of object- or scene-specific processing in the brain. However, their RSA correlation (Cichy et al., 2017) or encoding (Loke et al., 2025) curves were remarkably similar for both object- and scene-trained DCNN models; possibly because scene-trained DCNNs also learn to detect objects (Zhou et al., 2014). In our data, we also see that DCNNs trained with different objectives on different training datasets yield similar temporal encoding profiles (**Supplementary Fig. S4A+B**). However, when we vary the input to the pretrained models (by masking either scenes or objects) we obtain clearly distinct temporal profiles across many different DCNN backbones. This highlights that manipulating the inputs to the encoding model (i.e. the input to the already pre-trained DCNNs when extracting its features), rather than the DCNN training data, is a promising avenue for future studies interested in disentangling visual information processing in brain responses to natural images; indeed, a similar logic has recently been used to show that texture features are better predictors of EEG responses to natural images than isolated objects (Loke et al., 2026). Overall, our results add to a growing body of work showing that variance in EEG signals can be meaningfully linked to spatially-localized image features (Hansen et al., 2021; Müller et al., 2026), differentiating, for example, the image features driving EEG activation under different task instructions (Hansen et al., 2025).

### Potential confounds for temporal encoding differences

In our analyses, we addressed a number of factors that could have potentially confounded the temporal separation of scene and object information processing. First, we demonstrated that the observed temporal difference between the Objects and the Scene models are only present when applying the bounding boxes to the original locations, in the matching images. When instead shifting the bounding boxes away from their original locations or using the bounding boxes of a randomly chosen other image, the temporal differences disappear. These results confirm the distinct temporal profiles of object and scene processing.

Second, we showed that the temporal differences are not an artifact of the many additional sharp edges that are introduced in the images by masking pixels within the rectangular bounding boxes before feeding them to the DCNN feature extractor. The temporal differences remains when first extracting the DCNN features for the original images and only then applying the bounding boxes to the extracted features. However, in this latter approach the separation of object from scene information becomes less precise as DCNN features become spatially pooled in increasingly smaller feature maps compared to their input, and information at the boundaries between objects and the scene may blend into each other with increasing convolutional steps. Still, this approach strengthens our main findings by showing that in either case, a temporal distinction between object and scene processing consistently surfaces.

Third, we investigated whether the separation of object from scene information coincides with the spatial separation of features between fovea and periphery. In the present EEG dataset, we previously found a temporal hierarchy in neural processing of peripheral and foveal information, with peripheral encoding (around 100 ms after stimulus onset) preceding foveal encoding (around 150 ms), coinciding exactly with our peak encoding windows for scene and object information. Moreover, fMRI evidence suggests that fovea-periphery biases in human visual cortex relate to the functional specialization of category-selective regions„ such that foveally-biased cortex tends to overlap with face-selective regions, whereas peripherally-biased cortex overlaps scene-selective regions (Hasson et al., 2002; Arcaro and Livingstone, 2017), suggesting a close link between spatial sampling and specialized information processing (Groen et al., 2022). However, while objects do tend to occur more frequently at central locations in our image set, our control analysis showed that the object and scene encoding profiles are not driven by foveal vs. peripheral information sampling, respectively. This result is consistent with recent work showing that spatial biases do not fully account for fMRI response selectivity (Silson et al., 2024).

### Using image-grounded encoding to understand what defines objects and scenes

To construct our image-grounded encoding models, we needed an initial definition of what ‘counts’ as an object versus a background scene element. While it is surprisingly difficult to formally define what constitutes an object vs. a part of a scene (Groen et al., 2017), this distinction is often intuitive for human observers. This motivated us to use a top-down approach, starting with human-derived manual annotations of objects within natural scenes, and then using DCNNs to provide us with object- and scene-bound features for predicting neural signals.

Nevertheless, some of the labeled object classes are inherently ambiguous: they may be perceived as an object if it appears in one context, but as part of a scene in another. In our study, there are multiple such instances, including benches and doors. Importantly, we show that our image-grounded encoding approach allows for a data-driven identification of the perceived ‘objectness’ of a given object class (or scene element). Doors can serve as an illustrative example for this ambiguity, as they have many properties that would typically define an object: they can be moved individually compared to theirs frames; they can be interacted with (Bainbridge and Oliva, 2015); their spatial extent and also identity is defined most strongly by their shape (Landau et al., 1988; Cutzu and Tarr, 1997), not their texture (Geirhos et al., 2018), color or other surface level properties. Yet, they fell out of our individual object analysis as an outlier, suggesting that they are perceived more like part of the scene than individual objects (similarly to the more prominent and intuitive example of trees). Thus, while our encoding models necessitated an initial definition to separate of objects and scenes, the encoding model performances themselves can be used to adjust or refine such definitions based on their observed neural processing profiles.

Apart from definitions, an enduring question in the study of functional specialization in visual perception is which aspects of it are driven primarily by ‘low-level’ features (e.g. spatial frequency distributions, Rajimehr et al., 2011) versus ‘high-level’ information (e.g. semantic categories, Walther et al., 2009). Prior research in the temporal domain has examined different types of information processing during natural image perception by contrasting, for example, image-computable low-level image features with high-level, semantic object or scene labels (Greene and Hansen, 2020; Bartnik et al., 2025a). Our DCNN-based image-grounded encoding modeling deviates from these approaches: since our object- and scene-bound features are derived from the same DCNN backbone and the same network layers, they may be considered as being on ‘equal footing’ in terms of feature complexity, and we did not focus our analyses on uncovering whether different types of features drove scene-vs. object temporal encoding profiles (e.g. more low-level features for scene compared to object encoding).

However, these types of questions could in principle be addressed in an image-grounded encoding framework as well, for example by contrasting encoding performances across different model layers. An initial comparison of encoding for scene vs. object-bound features across different layers (**Supplementary Fig. S3**) indicates that both peak encoding and the temporal differences are most pronounced in early and layers, suggesting a primary role for low- or mid-level features in driving the observed temporal differences in encoding profiles. However, the same analysis shows that time-averaged encoding performance increases slightly from lower- to higher DCNN model layers; this shift appears to be driven by a comparatively stronger encoding of object information compared to scene information at later EEG time windows. Future studies are necessary to determine whether these patterns are statistically robust, and whether they hold over multiple DCNN model architectures and model depths.

### Resolving the trade-off between controllability and ecological validity

Typically, experiments targeting the neural mechanisms underlying perception face a trade-off between the controllability of experimental parameters versus their ecological validity. In their extremes, one excludes the other: a manually crafted visual stimulus, where each component (such as lines, circles, colors, etc.) is controlled by its own parameter, is highly unlikely to be found in nature - and these stimuli have drastically different statistical properties compared to natural stimuli (e.g., David et al., 2004). Contrarily, natural scenes are a complex collection of lines, colors, and other components, that are difficult to experimentally manipulate without losing image naturalness (although state-of-the-art diffusion models may offer a new solution, see e.g., Garcia Cerdas et al., 2025). Here, we propose that instead of considering individual stimuli as separate experimental conditions, one can use image-grounded encoding models that each receive different information from the same stimuli as the experimental conditions. By using pixel-level annotations to manipulate the model inputs, rather than the stimuli presented to participants, we recover a form of experimental control thought to be lost when using natural stimuli.

This method could be extended to experiments in virtual environments, further increasing the experimental control over visual information whilst increasing the level of immersion in the natural stimuli. This approach can also be employed in task-based paradigms, e.g. when manipulating covert attention using spatial cues or object-based priming, to identify how the encoding of the same natural, visual information changes under task influences. Finally, image-grounded encoding models can easily be applied to other brain measurement modalities (e.g., fMRI) to identify functional specialization, and expanded to multimodal sensory stimulation (e.g., audiovisual).

### Limitations and suggestions for future work

First, our stimuli exclusively consisted of Amsterdam street scenes, with limited diversity in object classes and scene parts. For example, there were few water-labeled pixels and encoding contribution of those pixels was near-zero. Future research could determine whether these classes have a stronger encoding contribution in more diverse images depicting larger regions of those classes. Second, the indoor-detection task may have influenced the image features participants attended to — e.g., prioritizing information that helps differentiate indoor from outdoor scenes. However, since the object classes and scene parts we analyzed are both primarily found in outdoor scenes, we expect any task influences to similarly affect their processing. Furthermore, our prior work suggests limited influence of task instructions on early EEG responses to natural scenes (Groen et al., 2016; Bartnik et al., 2025a). Still, future research should validate if the same temporal differences occur across multiple tasks instructions and investigate how tasks and attention interact with the temporal distinction between object and scene processing.

Further, it remains unclear where in visual cortex the temporal differences between object and scene processing originate. One possibility is that the EEG signal reflects neural activity of two separate regions specialized in scene and object processing (e.g., parahippocampal place area (PPA) vs. lateral occipital complex (LOC)). Another possibility is that it reflects the activity of the same region that processes object and scene information on a different time scale likely due to different low-level visual features, such as the primary visual cortex (V1). Prior work used intracranial EEG (iEEG) to study the joint spatio-temporal dynamics of object and scene processing and found that scene-selective electrodes within scene-selective regions respond earlier to scenes than object-selective electrodes within the same regions, while this difference is absent in more object-selective regions such as the fusiform gyrus and the lateral temporal complex (Vlcek et al., 2020). Kaiser et al. (2020) instead combined fMRI with EEG recordings and found that early discriminability of scenes seems to be rooted in V1, while the discriminability at time points later than 250 ms seems to originate in higher-level visual cortical regions such as PPA and the occipital place area (OPA). Bartnik et al. (2025a) showcased how fMRI-EEG fusion can help elucidate the spatial origins of temporal signatures of the processing of specific types of visual information, however they only compared different scene-selective regions. Together, these studies indicate that temporal differences of object and scene processing in EEG signals could be a joined signature of a spatial segregation of specialized processing and the processing of different features in early visual cortex. Future studies could use fMRI-EEG fusion to attempt to localize the sources of temporal profiles of object and scene processing.

## Conclusion

We used deep neural network-based encoding models in combination with image annotations to delineate temporal dynamics of object- and scene-specific processing in EEG responses to ecologically-valid, natural scene images. We show that object encoding shows a delayed temporal profile compared to scene encoding and identify the unique contribution of individual object classes and scene elements to these profiles. These results suggest that object and scene processing in the human brain differentially unfold over time, in addition to being routed towards distinct brain regions. Our image-grounded encoding modeling approach opens up new possibilities for disentangling how individual components of the real-world visual environments are encoded in the human brain.

## Supplementary Materials

**Figure S1:**
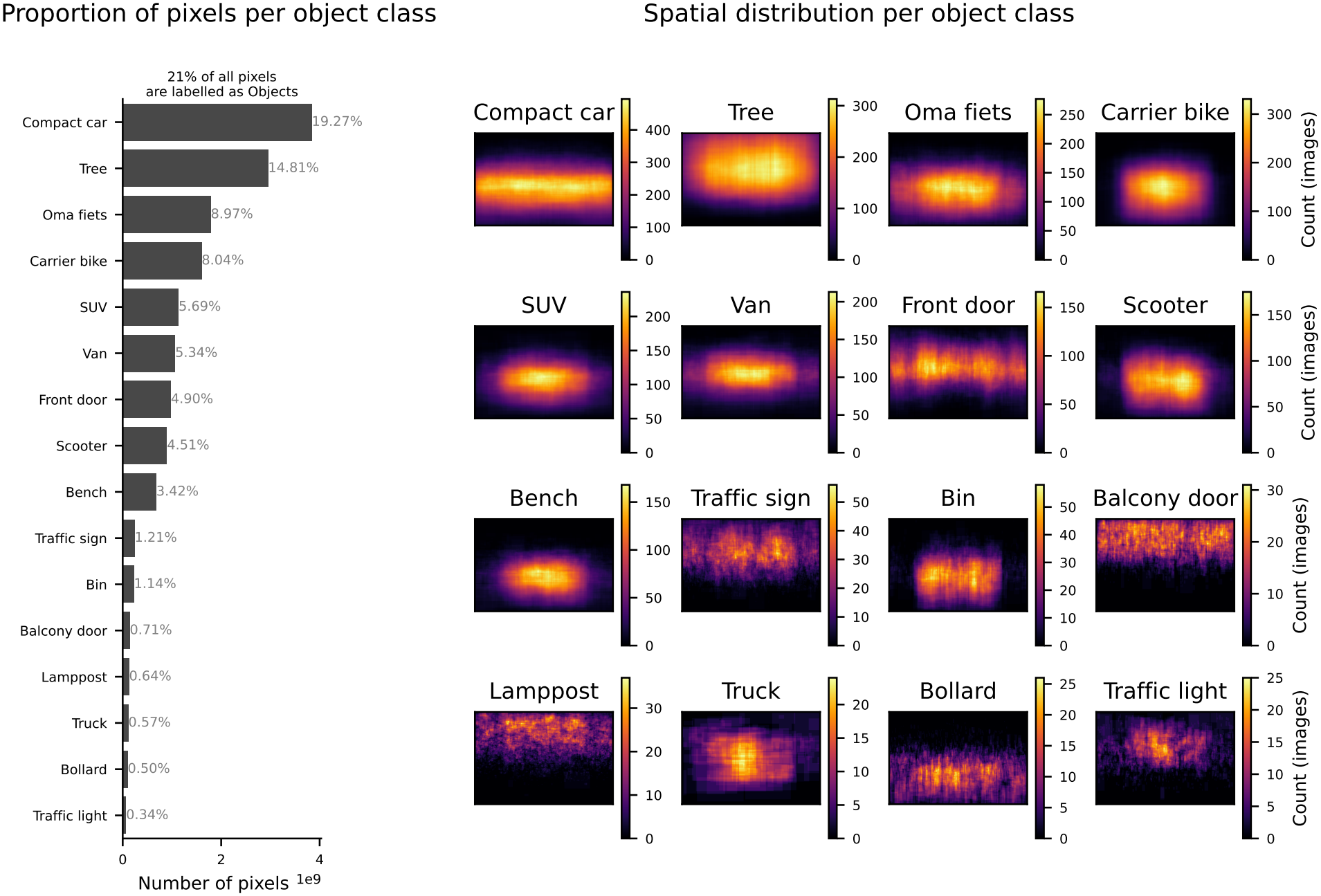
Distribution of annotated object pixels. Barplot shows the number of total pixels per object class, summed across all images. Heatmaps show the spatial distribution of the counts of annotated pixels per object class.

**Figure S2:**
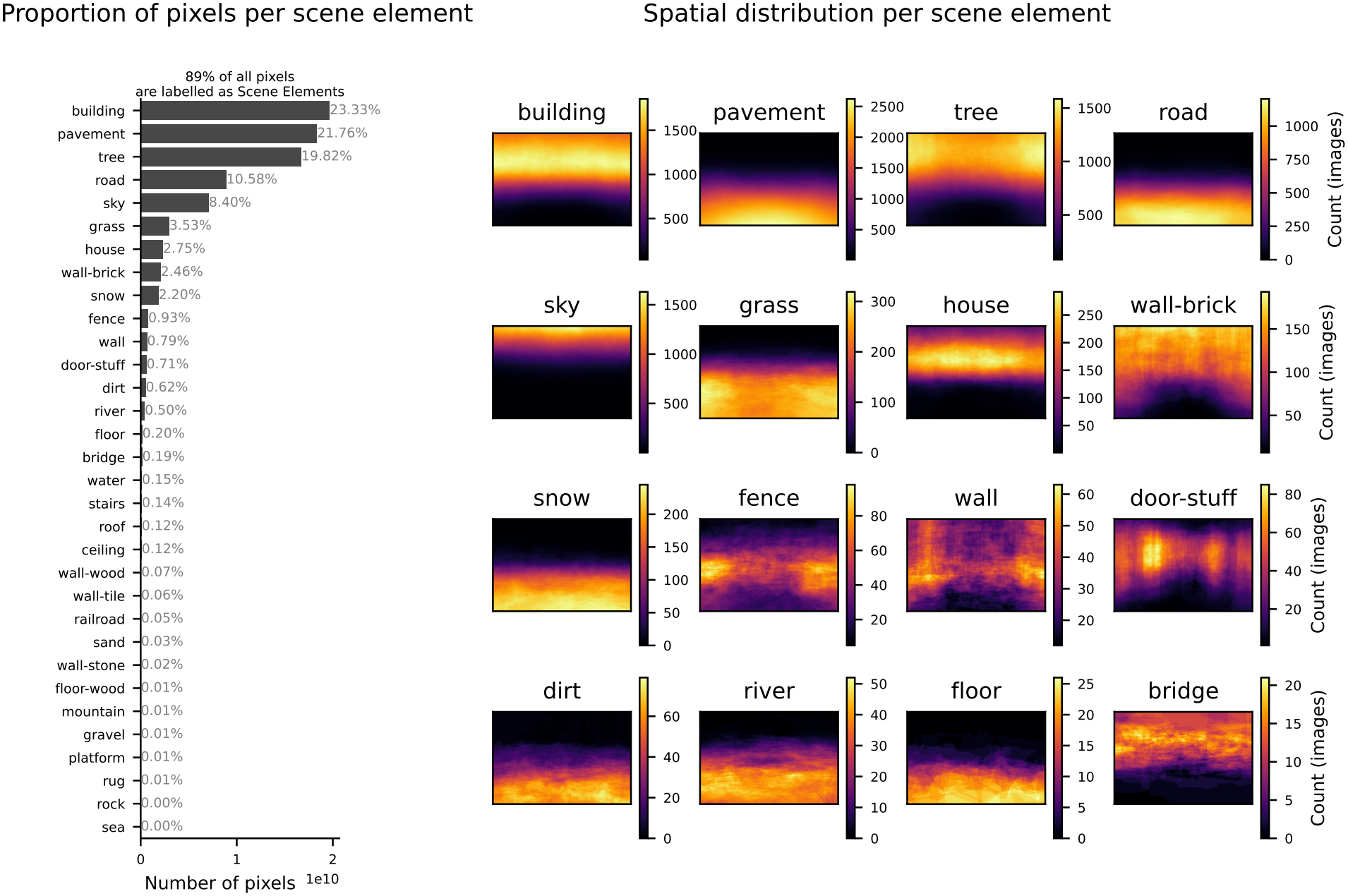
Distribution of annotated scene element pixels. Barplot shows the number of total pixels per scene element, summed across all images. Heatmaps show the spatial distribution of the counts of annotated pixels per scene element.

**Figure S3:**
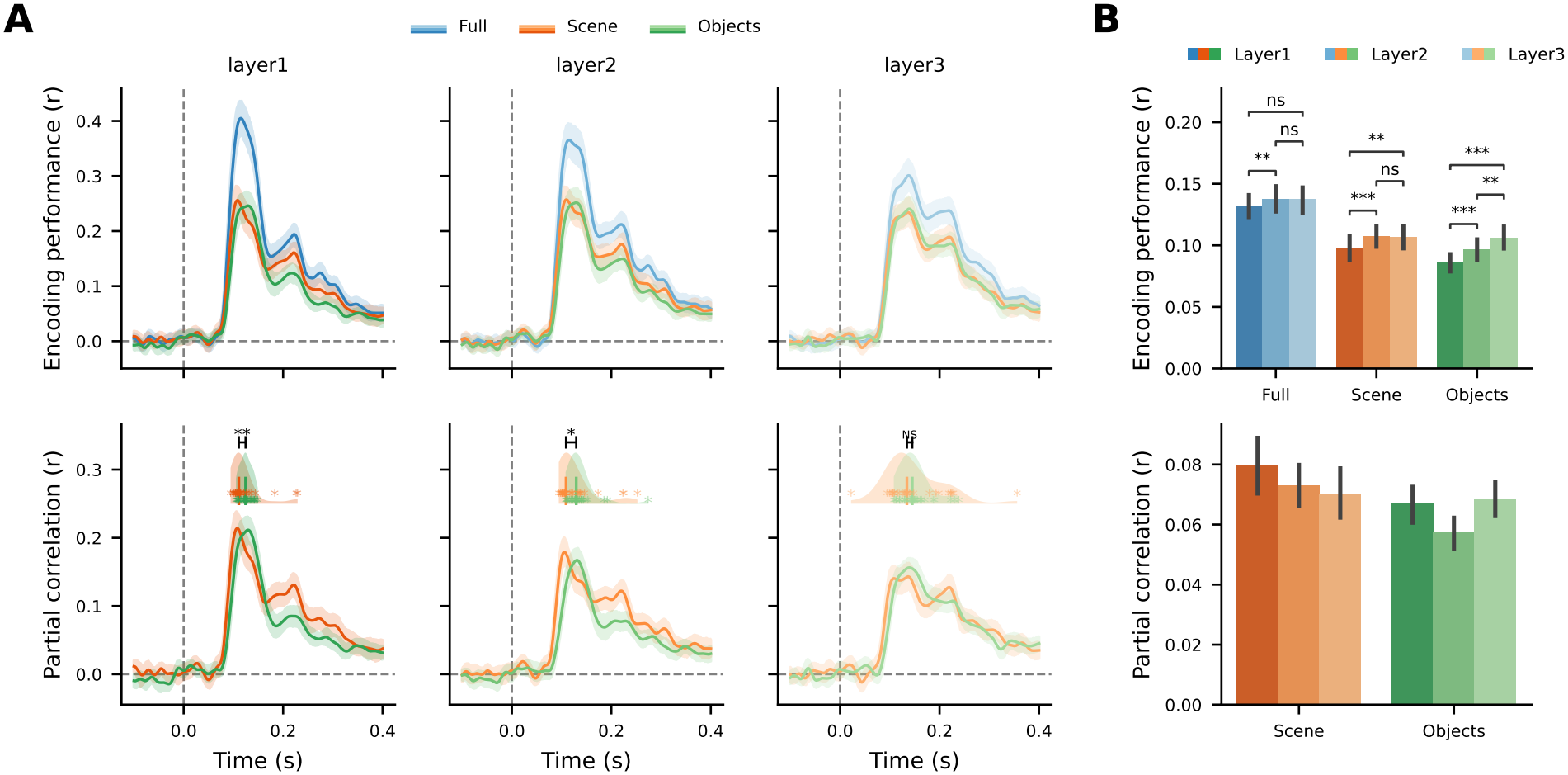
Encoding performance of individual DCNN layers. **A** Average encoding performance across participants when using different layers of AlexNet, for the correlation of each of the three main encoding models (Full, Scene, Objects; top row) and partial correlation between the Scene and Objects models (bottom row). **B** Same as in A, but for correlations (top) and partial correlations (bottom) averaged over time (t>0).

**Figure S4:**
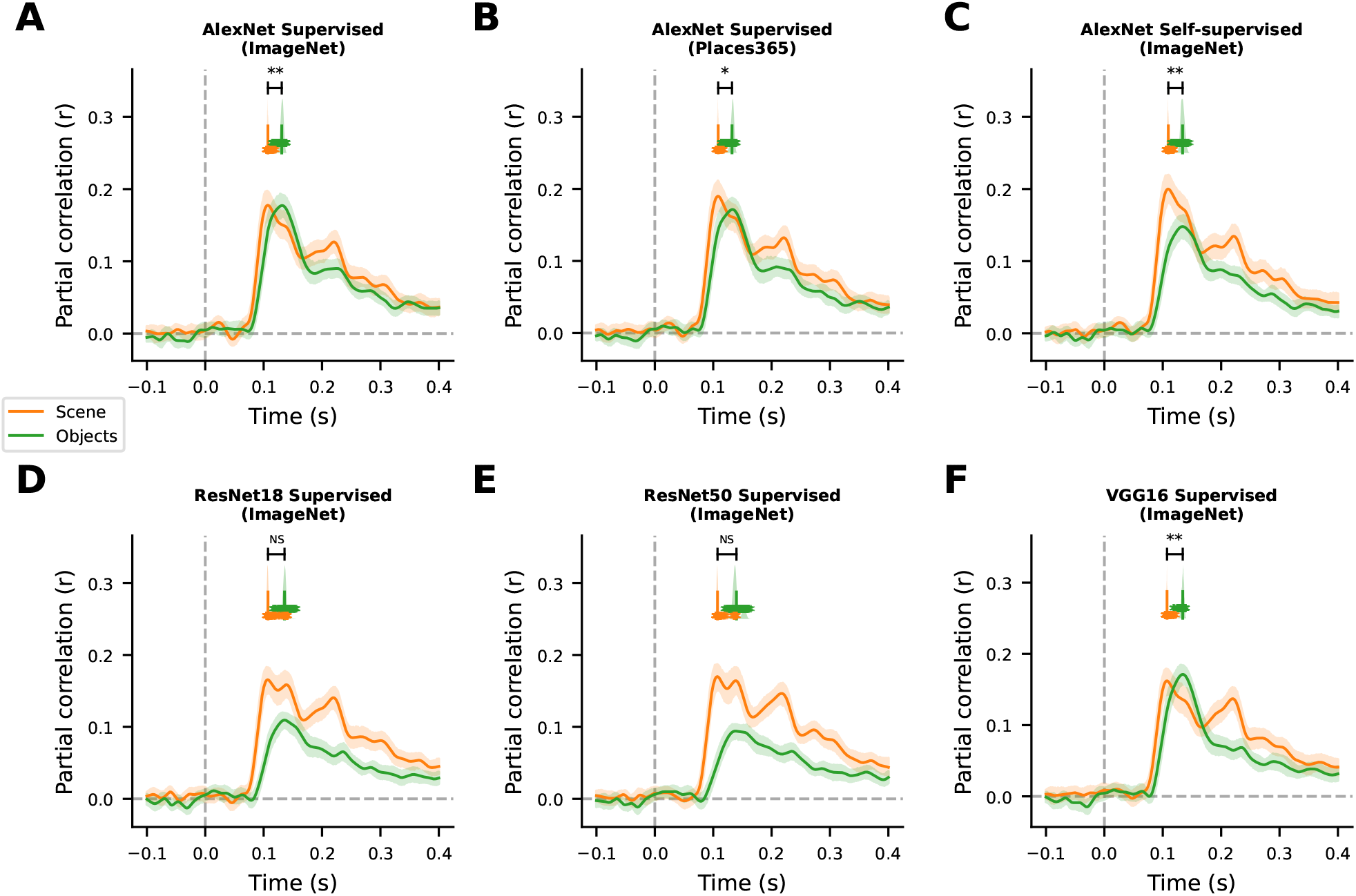
Temporal difference between scene and object encoding remains robust for all training regimes and some architectures. Each panel shows the average encoding performances of the Objects and Scene encoding models based on the features of different CNNs across participants over time Top row shows results from different training regimes and bottom row shows results from different CNN architectures. Coloured asterisks and distributions show the time points of maximum encoding performance per bootstrap iteration, vertical bars indicate the means across iterations and black asterisks above indicate statistical differences between the respective Objects and Scene models. **A** AlexNet trained on object-classification (supervised) on ImageNet. **B** AlexNet trained on scene-classification (supervised) on Places365. **C** AlexNet trained using contrastive learning (self-supervised) on ImageNet. **D** ResNet18 trained on object-classification (supervised) on ImageNet. **E** ResNet50 trained on object-classification (supervised) on ImageNet. **F** VGG16 trained on object-classification (supervised) on ImageNet.

**Figure S5:**
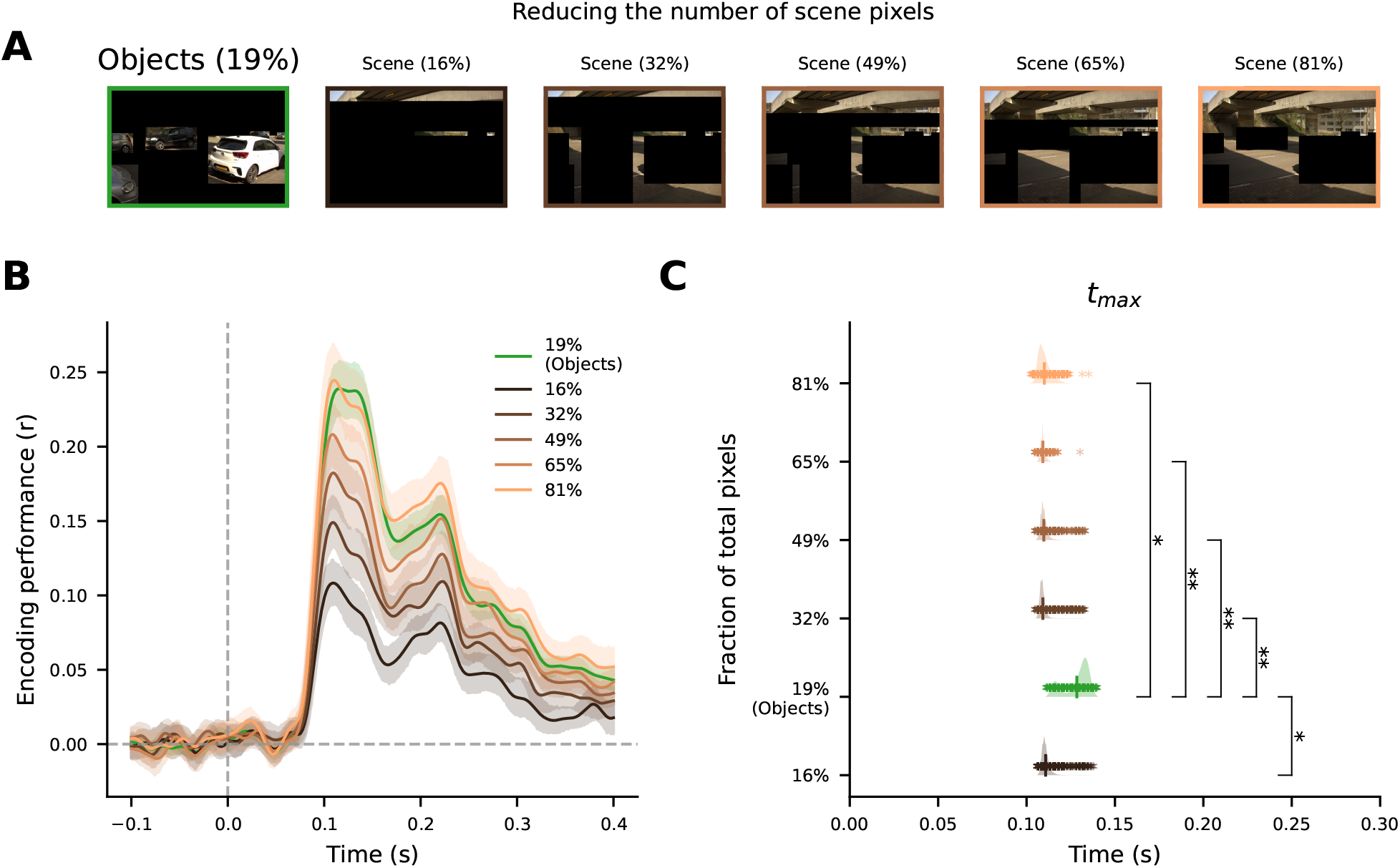
Temporal difference between scene and object encoding remains robust when matching the number of scene and object pixels. **A)** Example image with the Objects version (left) and multiple Scene versions in which the number of non-masked scene pixels was systematically decreased; the percentage above each stimulus refers to the total proportion of pixels that were left intact (from right to left: 100% of scene pixels, resulting in a total proportion of intact pixels of 81%; 80% of scene -> 65% of total; 60% of scene -> 49% of total; 40% of scene -> 32% of total; 20% of scene -> 16% of total; Object version has 19% of the total pixels). **B)** Average encoding performance across participants over time for the Objects model and versions of the Scene model receiving a reduced proportion of scene pixels. **C)** Distribution of the timepoints of maximum encoding performance for each model shown in B, vertically ordered by the total fraction of intact pixels. Coloured asterisks and distributions show the time points of maximum encoding performance per bootstrap iteration, vertical bars indicate the means across iterations and black asterisks on the indicate statistical differences of each pairwise comparison between the Scene model versions and the Objects model.

**Figure S6:**
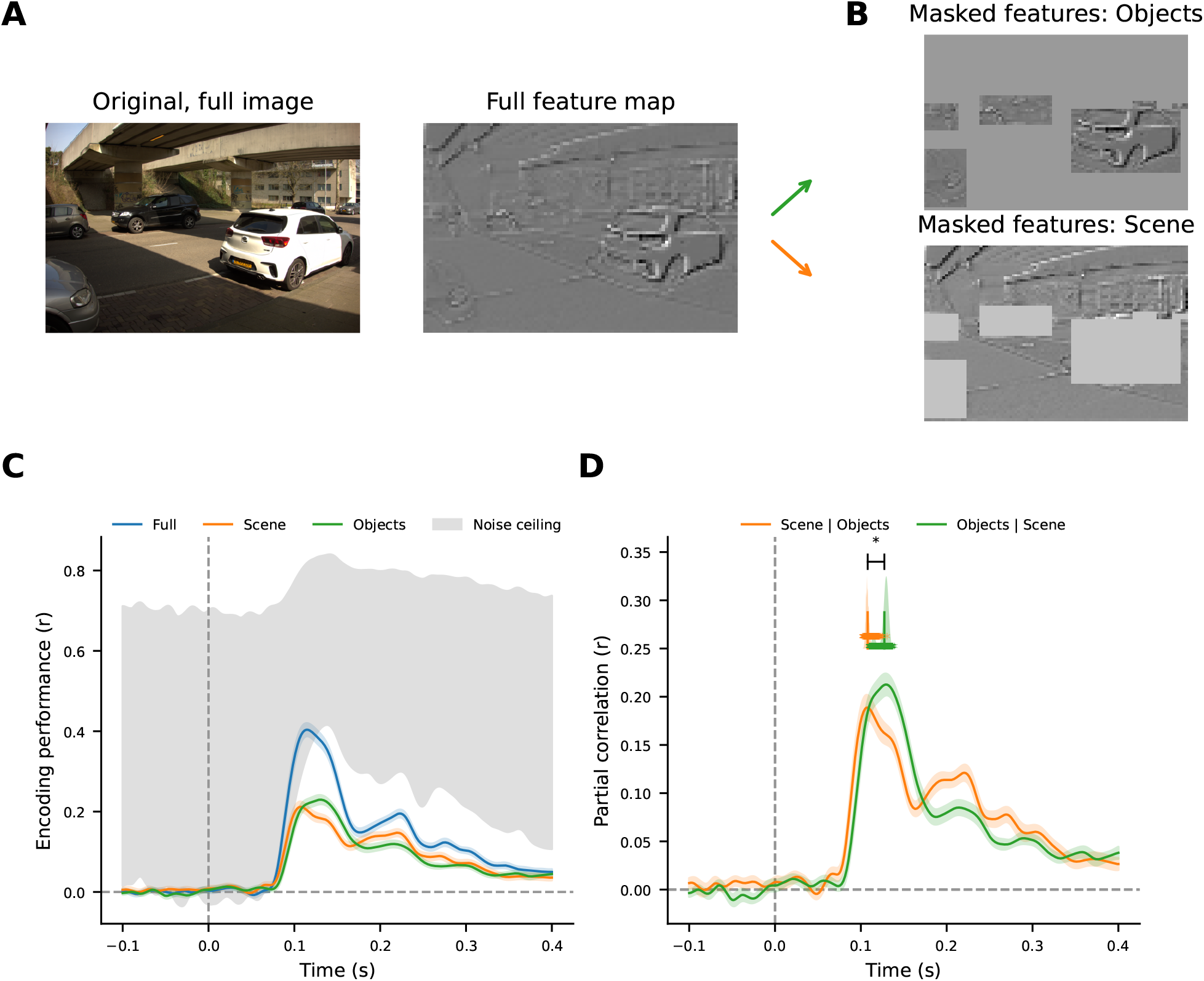
Temporal difference between scene and object encoding remains robust when bounding boxes are applied to features of intact image. **A** Example image that is fed to the CNN in its full, intact version, for which the full, intact CNN feature maps are extracted. The object annotations are then applied to these feature maps, to create the Objects and the Scene encoding model inputs, which can be seen in **B). C** Left panel shows the average encoding performance across participants over time for models built on the masked features shown in B) instead of the masking already being applied on the input images to the CNN. Shaded gray areas show the upper and lower bound of the estimated noise ceiling computed across stimulus repetitions on the test set per participant. Right panel shows the average partial correlation between the Objects and the Scene models across participants over time. Coloured asterisks and distributions show the time points of maximum encoding performance per bootstrap iteration, vertical bars indicate the means across iterations and black asterisks above indicate statistical differences.

**Figure S7:**
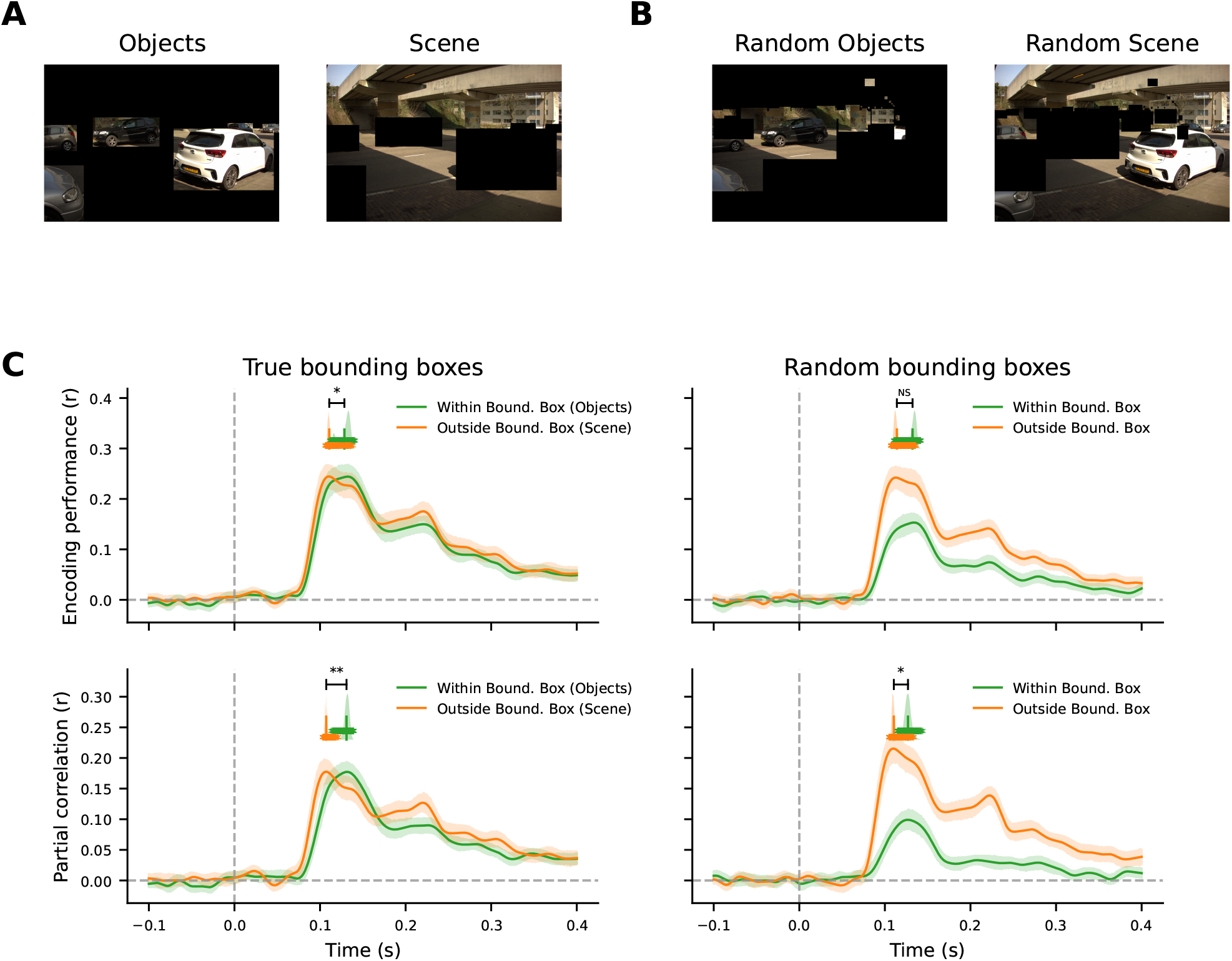
Temporal difference between scene and object encoding is disappears when using random bounding boxes. **A)** Example image with the true bounding boxes applied to get the Objects version (left) and the Scene version (right) **B)** Same as in A), but for applying the bounding boxes from a randomly chosen image. **C)** Average encoding performance (top) and partial correlation (bottom) across participants over time for models receiving mask inputs (pixels within bounding boxes kept intact in green; pixels outside bounding boxes kept intact in orange) when the true bounding boxes per image were used (left) or when the bounding boxes were randomly assigned to another image (right).

**Figure S8:**
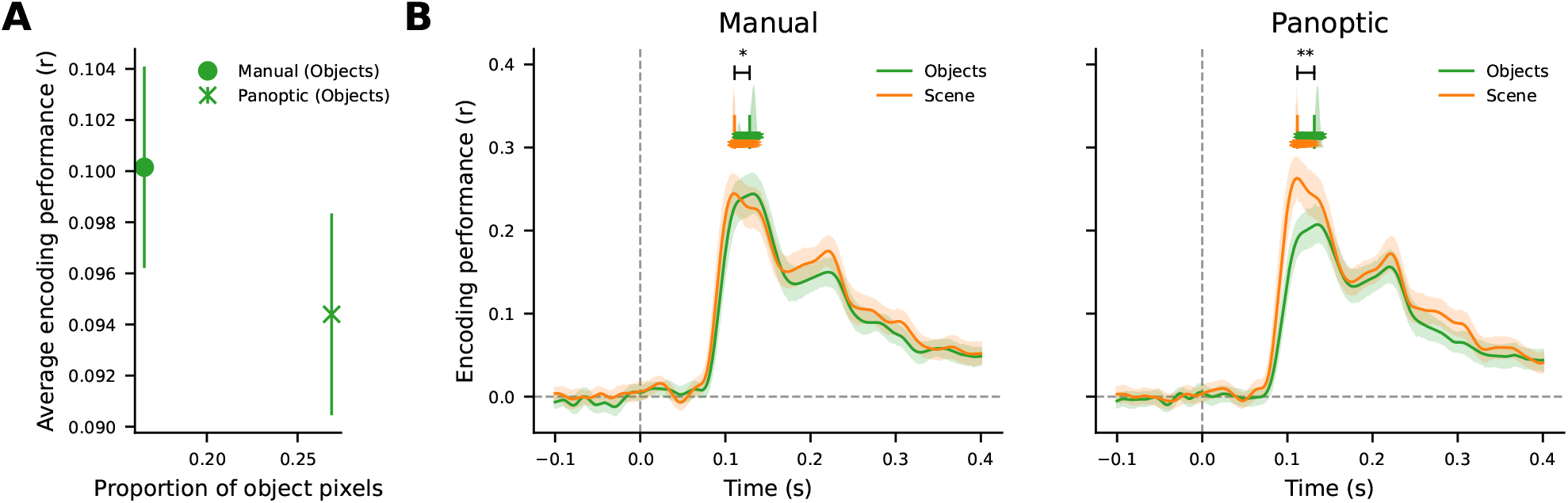
Temporal difference between scene and object encoding remains robust when using automated annotations. **A)** Average encoding performance across participants for models using the manual OADS object annotations or a selection of the panoptic image segmentation that correspond to a similar set of objects as used in OADS, averaged across all time point after stimulus onset. **B)** Time-resolved encoding performances averaged across participants for the same Objects models as in A) together with their corresponding Scene model versions, for the manual annotations (left; same data as shown in Fig. 2B) and the panoptic annotations (right). Coloured asterisks and distributions show the time points of maximum encoding performance per bootstrap iteration, vertical bars indicate the means across iterations and black asterisks above indicate statistical differences.

## Data availability

The stimuli, EEG data and encoding models results that support the findings of this study are available on OSF (https://osf.io/cmw4a) with the identifier “DOI 10.17605/OSF.IO/TU34H”. Stimuli are available with blurred faces and license plates to preserve privacy. Due to privacy constraints, original (non-blurred) stimuli can be requested on a case-by-case basis.

## Code availability

The code used to build encoding models, perform statistical tests and produce figures can be found on github: https://github.com/niklas-mueller/oads_eeg_object_scene_encoding

## Notes

https://osf.io/cmw4a

### Competing Interest Statement

The authors have declared no competing interest.

### Summary of Updates

Figure 3+4 revised; Supplementary figures updates

